# Migration, acculturation, and the maintenance of between-group cultural variation

**DOI:** 10.1101/234807

**Authors:** Alex Mesoudi

## Abstract

How do migration and acculturation (i.e. psychological or behavioral change resulting from migration) affect within- and between-group cultural variation? Here I answer this question by drawing analogies between genetic and cultural evolution. Population genetic models show that migration rapidly breaks down between-group genetic structure. In cultural evolution, however, migrants or their descendants can acculturate to local behaviors via social learning processes such as conformity, potentially preventing migration from eliminating between-group cultural variation. An analysis of the empirical literature on migration suggests that acculturation is common, with second and subsequent migrant generations shifting, sometimes substantially, towards the cultural values of the adopted society. Yet there is little understanding of the individual-level dynamics that underlie these population-level shifts. To explore this formally, I present models quantifying the effect of migration and acculturation on between-group cultural variation, for both neutral and costly cooperative traits. In the models, between-group cultural variation, measured using F statistics, is eliminated by migration and maintained by conformist acculturation. The extent of acculturation is determined by the strength of conformist bias and the number of demonstrators from whom individuals learn. Acculturation is countered by assortation, the tendency for individuals to preferentially interact with culturally-similar others. Unlike neutral traits, cooperative traits can additionally be maintained by payoff-biased social learning, but only in the presence of strong sanctioning institutions. Overall, the models show that surprisingly little conformist acculturation is required to maintain realistic amounts of between-group cultural diversity. While these models provide insight into the potential dynamics of acculturation and migration in cultural evolution, they also highlight the need for more empirical research into the individual-level learning biases that underlie migrant acculturation.

## Introduction

Humans exhibit extensive between-group cultural variation [1–3] as documented by anthropologists [4], psychologists [5] and economists [6] in domains such as languages, customs, marriage practices, cooperative norms, religious beliefs and social organisation. While such traits certainly vary within societies, they typically characterize, and sometimes even define, entire social groups. At the same time, migration has been a constant fixture of our species for millenia [7]. *Homo sapiens* migrated out of Africa in multiple waves from 100kya [8], and migration continued within and between global areas throughout prehistory [9,10]. In the modern era, improved transportation technologies such as steamships allowed mass migration to transform the USA in the early 20th century [11], while international migration has increased in volume (number of migrants) and scope (diversity of destinations) in the last century [12].

In a sense, these two phenomena - extensive between-group cultural variation and frequent migration - are contradictory. It is well known in population genetics that even small amounts of migration can rapidly break down between-group genetic structure [13,14]. If migration acts on cultural variation in the same way as it acts on genetic variation, we would expect that our species’ frequent migration would rapidly destroy between-group cultural variation, resulting in a homogenous mass world culture. At the least, we would expect similar levels of between-group cultural variation as between-group genetic evolution, yet the former is more than an order of magnitude greater than the latter [3]. However, cultural evolution does not act in the same way as genetic evolution. Migrants are stuck with their genes, but they may change their cultural traits, and often do so within one or two generations. This process of *acculturation*, defined as psychological or behavioral change resulting from migration [15], could, in principle, maintain between-group cultural variation even in the face of frequent migration.

Here I adapt population genetic models of migration to the cultural case to ask: (i) how strong does acculturation need to be, and what form should it take, in order to maintain different amounts of between-group cultural variation in the face of different levels of migration? (ii) are these levels of migration, acculturation and between-group cultural structure empirically plausible? I follow previous cultural evolution researchers [16,17] in adapting population genetic methods and concepts to analyse cultural change, given that genetic and cultural change are both systems of inherited variation [18]. While past models examined the effect of migration on cultural diversity [16] and the effect of acculturation in specific communities [19] and on multiculturalism [20], no past research has explicitly linked migration and acculturation to the maintenance of between-group cultural variation using quantitative measures of the latter. And while migration is often studied in sociology [21], psychology [15] and cultural anthropology [22], much of this work focuses on the proximate level (e.g. the lived experiences of migrants) and does not formally link individual-level migration and acculturation processes to population-level cultural dynamics.

These questions have relevance beyond academia. Migration and acculturation have always been major factors in political and public discourse, and never more so than in today’s increasingly interconnected world. As noted above, economic migration is at record global levels [12]. The involuntary migration of refugees has also increased. The ‘European migrant crisis’ has seen unprecedented numbers of people crossing the Mediterranean from Syria and other conflict areas [23]. As migration increases, public opposition to immigration often also rises [24]. Much public opposition reflects beliefs that immigrants are incapable of acculturating, and that migration will consequently weaken or destroy existing cultural traditions in receiving countries. In the 2013 British Social Attitudes survey [25], 77% of respondents wanted immigration levels reduced, 45% said immigrants undermined Britain’s cultural life (versus 35% who said they enriched it), and half agreed that one cannot be “truly British” unless one has British ancestry (51% agree) and shares British customs and values (50% agree). Politicians often appeal to such fears surrounding supposedly non-acculturating immigrants. Marine Le Pen of the French *Front national* has said “Immigration is an organized replacement of our population. This threatens our very survival. We don’t have the means to integrate those who are already here.” [26]. Nigel Farage of the *UK Independence Party* has said “If we went to virtually every town up eastern England and spoke to people about how they felt their town or city had changed for the last 10 to 15 years, there is a deep level of discomfort. Because if you have immigration at these sort of levels, integration doesn’t happen” [27]. My aim here is not to take a stance on the normative and political issue of migration policy, only to scrutinize the logical and empirical claims inherent in such statements and policies, e.g. that immigrants can never acculturate (Le Pen), or that acculturation depends on the level of migration (Farage).

After reviewing the evidence for migrant acculturation, I then justify the use of conformist social learning as a candidate for an acculturation mechanism and explain how the present study differs from previous models of conformity and migration. Model 1 then explores how migration and conformist acculturation interact to maintain, reduce or eliminate between-group cultural variation in functionally-neutral cultural traits. Model 2 examines how migration and acculturation affect a non-neutral cooperative trait that is individually costly but group beneficial. Models presented are recursion-based simulations tracking the frequency of cultural traits over time, written in R [28] with code available at https://github.com/amesoudi/migrationmodels.

### Evidence for migrant acculturation

There is a growing literature in economics, sociology and psychology that compares quantitative measures of behavioral or psychological traits across multiple generations of immigrants. Psychologists have tested migrants of non-Western heritage living in Western countries on measures that exhibit cross-cultural variation, such as collectivism, social attribution, self-enhancement and self-esteem [29–31]. Economists and political scientists have measured traits such as institutional trust via large-scale surveys in multiple generations of migrants [32–34]. This work aims to determine whether immigrants (or their descendants) retain the trait values of their country of birth or shift towards the trait values of their adopted country, in those cases where such values are different. If there is a shift, studies often determine its magnitude, how it varies with different combinations of adopted and birth countries, how it varies with age of migration (for first generation migrants), and how it persists over subsequent generations (second, third etc. generations). Using standard terminology, first generation migrants are defined as those born and raised in one country and who moved to another country after the age of 14, second generation migrants are children of first generation migrants, third generation are children of second generation, and so on.

Table S1 and Figure S1 present some example findings from the economics, sociology and psychology literatures, where mean values from multiple migrant generations are reported. It is necessarily selective, but broadly representative of other studies [19,33,34]. From Table S1, and those other studies, we can make the following empirical generalizations which inform the models in the current study:

- **Complete one-generation assimilation is rare.** It does not appear that first generation migrants typically acculturate/assimilate completely to their adopted country’s values. Table S1 shows several traits for which first generation migrants differ from non-migrants, often substantially, and in the direction predicted by the cross-cultural research mentioned above [2,6]. For example, one study [29] found that first generation British Bangladeshi migrants are more collectivistic and show more situational, and less dispositional, social attribution compared to White British non-migrants. This reflects findings in cross-cultural psychology that Western societies are less collectivistic, and show more dispositional and less situational attribution, than non-Western societies [35]. Another study [32] found that non-Western first generation migrants to high trust countries such as Denmark, Norway and Sweden show much less trust, again in line with existing cross-cultural research. While there may be other traits not shown in Table S1 that do show complete within-generation assimilation, this seems rare. A wealth of literature shows that cultural history is important, and people are shaped by the values they acquire during childhood and early life [33,34].
- **Acculturation is common and occurs over multiple generations.** While no trait in Table S1 shows complete, immediate, one-generation assimilation, most traits show some degree of gradual acculturation. First generation migrants often show a partial shift towards their adopted country’s values, and second and further generations show shifts that are as large or larger. In many cases this shift is quite substantial, with second generation migrants sometimes moving more than 50% of the way towards non-migrant values from their first generation parents’ or heritage country’s values (see Figure S1). For example, 2nd generation British Bangladeshis have shifted around 50% towards local non-migrant values of collectivism and social attribution from their 1st generation parents’ values [29]. In other cases shifts are only evident after multiple generations. In a rare study of four generations of migrants living in the USA [33], 13 of 26 attitudes related to family, cooperation, morality, religion and government shifted of 50% or more towards adopted country values between the first and second generation, and 23 of 26 had shifted 50% or more by the fourth generation.
- **Acculturation rates vary across traits.** In the same population, some traits can shift substantially while others show little change. For example, there is little difference between first and second generation British Bangladeshis in religiosity and family contact, but a substantial shift in psychological traits such as collectivism or attributional style [29]. Similarly, acculturation rates in US-born Tongan Americans (i.e. second generation migrants) were estimated at approximately zero for some traits (e.g. sibling adoption) and as high as 0.87 for others (e.g. ranking of family lines) [19], while another study [33] found that trust-related attitudes show stronger acculturation than family and moral values.

While this empirical literature confirms that acculturation occurs, and provides estimates of population-level acculturation rates, they do not examine the individual-level dynamics (i.e. how migrants learn from others) that drive these population-level acculturation patterns, nor do they extrapolate from observed acculturation rates to long-term impacts on between-group cultural variation, as addressed below.

### Acculturation as conformist social learning

While acculturation as a population-level phenomenon certainly exists, there is no empirical research on the individual-level mechanisms by which an individual migrant alters their beliefs or practices as a result of learning from the wider community. The most plausible mechanism already modeled and tested in the cultural evolution literature is conformity [17,36–38], defined as an increased tendency to adopt the most common trait among a sample of other individuals (‘demonstrators’) relative to copying randomly. For example, if 70% of demonstrators are observed possessing trait A, then a conformist learner adopts trait A with a probability greater than 70%. An unbiased, non-conformist learner would adopt trait A with probability exactly 70%. Evolutionary models show that conformity is adaptive and thus likely to evolve in spatially variable environments connected by migration, and when individuals must choose between more than two traits [39], both features of the models implemented below. Both experimental [36,38,40] and real-world [41] evidence suggests that people are conformist. Conformity is also found in non-human species when naive individuals must acquire equally-rewarded cultural traits in a spatially heterogeneous environment with migration [42], again resembling the models used below, suggesting the general adaptiveness of a conformist social learning strategy.

Here I assume that conformity has already evolved (either genetically or culturally [43]) and explore its consequences for shaping between- and within-group cultural variation. Boyd and Richerson [17] and Henrich and Boyd [37] have previously shown that conformity can maintain between-group cultural variation. However, there are several limitations of those models for our understanding of conformity as an acculturation mechanism. First, there was no explicit measure of between-group cultural variation against which to compare model output to empirical data. No previous study has therefore asked or shown *how much* conformist acculturation is needed to maintain empirically plausible amounts of between-group cultural variation in the face of different levels of migration. Second, those models only featured two cultural traits and three demonstrators, which limits the scope for between-group cultural variation and the potency of conformity respectively. Subsequent models [39] have examined more than two traits, but using a different implementation with a conformity parameter that has an unclear individual level interpretation (see S1 Methods). Third, conformist social learning in those models occurs before migration and so does not affect new migrants. We are interested instead in how migrants potentially acculturate immediately after entering a new population. Finally, I extend such models to the maintenance of cooperative cultural traits maintained by punishment, to compare how acculturation acts to maintain between-group variation in neutral and cooperative traits. This is particularly important given that many debates surrounding the impact of immigration focus on migrant’s contributions to public goods and obeyance of cooperative social norms.

The novelty of this study therefore lies in:

- linking conformity to a quantitative measure of between-group variation, *F_ST_*, which has been used in the empirical literature, to ask *how much* conformist acculturation is needed to maintain plausible values of this measure in the face of migration, going beyond previous studies which have simply noted that conformity can maintain between-group variation
- extending Boyd and Richerson’s [17] model of conformity to more than two cultural traits and more than three demonstrators, including cases of joint-majority traits that emerge in such cases, using an implementation that has an empirically interpretable conformity parameter
- implementing conformist acculturation as occurring after migration, not before, such that new migrants can potentially acculturate
- directly comparing neutral and cooperative traits on the above measures, given the focus of much acculturation research on cooperation-related cultural values such as trust and corruption

## Model 1: Migration, acculturation and cultural diversity

I start with a simple model of migration borrowed from population genetics, Wright’s island model [13,14], to which I later add acculturation. Parameters are listed in Table 1. Assume a large population divided into *s* equally sized and partially isolated sub-populations. Consider a trait (originally an allele, but here a cultural trait) that has mean frequency across the entire population of 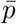. Every timestep, each individual migrates with probability *m*. Migration is random and simultaneous, such that a proportion *m* of individuals are randomly removed and this pool of migrants is randomly allocated across all newly-vacant spots in the entire population. Hence a proportion *m* of individuals in each sub-population will be immigrants, taken from a pool with trait frequency 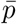.

**Table 1:** Parameters in Model 1

| Parameter | Definition |
| --- | --- |
| $s$ | The number of sub-populations, and also the number of cultural traits. |
| $p$ | The frequency of a trait in a sub-population |
| $m$ | Migration rate: the probability in one timestep that an individual moves to a randomly chosen sub-population. Equivalently, the proportion of the population that moves in one timestep. |
| $n$ | The number of demonstrators from whom individuals learn during acculturation. |
| $a$ | Acculturation rate: the strength of conformist social learning, defined as the degree to which an individual is more likely to adopt the most-common trait among $n$ demonstrators chosen from their sub-population, compared to unbiased, random copying. |
| $r$ | Assortation strength. With probability $1 - r$, the $n$ demonstrators are chosen at random. With probability $r$, sets of demonstrators all have the same cultural trait. |

Consider a particular sub-population in which the frequency of the trait at time *t* is *p*. For a randomly chosen trait in this sub-population, the trait either came from a non-migrant in the previous timestep with probability 1 − *m*, with the frequency among those non-migrants *p*_*t*−1_, or from a migrant with probability *m*, with the frequency among those migrants being 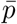. The overall frequency is therefore [13,14]:

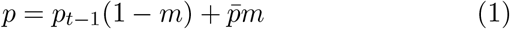

We are interested in how migration breaks down, and acculturation prevents the breakdown, of between-group cultural variation. Assume therefore that initially there is complete between-group cultural variation, and no within-group variation. Consequently, assume *s* cultural traits, and at time *t* = 0 all individuals in sub-population 1 have trait 1, all individuals in sub-population 2 have trait 2, and so on. To quantify the degree of population structure, I use Wright’s *F_ST_*, a measure typically used to quantify the degree of between-group genetic variation [13,44], but recently used to quantify population structure in cultural traits [3]. As a statistical measure, *F_ST_* is neutral to transmission mechanism, and simply measures the probability that two randomly-chosen individuals from the same sub-population carry the same trait, compared to two randomly-chosen individuals from the entire population (see S1 Methods). An *F_ST_* of 1 indicates complete between-group variation and no within-group variation. An *F_ST_* of 0 indicates no between-group variation, with all variation occurring within groups. Here, at time *t* = 0, *F_ST_* = 1, and we are interested in when this is maintained and when it drops.

It is difficult to obtain reliable real-world estimates of cultural *F_ST_* given the lack of quantitative data in fields such as cultural anthropology. Moreover, while group differences are often reported [4], within-population variation is seldom measured. Cultural evolution researchers have begun to address this, finding a mean cultural *F_ST_* of 0.08 for World Values Survey items [3] and 0.09 for a folktale [45]. Another study lists *F_ST_* values from 0.008 to 0.612 for various attitudes [46]. Clearly between-group cultural variation can vary greatly per trait and population(s), but *F_ST_* = 0.1 is perhaps a reasonable benchmark.

What is a plausible value of *m*? Migration rates differ across time periods and geographical scales. If a timestep in the model is taken as a biological generation, as it typically is in population genetics, then *m* is the probability that an individual migrates from their natal community during their lifetime. Lifetime migration rates have been estimated for several human populations within nation states (e.g. !Kung villages, or villages in Oxfordshire) at 0.1-0.33 [47]. Another interpretation of *m* is the proportion of the population that are migrants. In the UK this has increased from 0.042 in 1951 [48] to 0.133 in 2015 [49]. The USA has similar figures of 0.069 in 1950 and 0.134 in 2015 [50]. Some countries, such as Switzerland and Australia, reach a foreign-born proportion of the population close to 0.3 [51]. Representative low, typical and high values of *m* might therefore be 0.01, 0.1 and 0.3 respectively.

Following migration there is acculturation. As justified above, acculturation is implemented here as conformist social learning. Conformist acculturation is added to the island model in each timestep following the completion of all migration. It is assumed to apply to all individuals, not just new migrants. This allows acculturation to occur over multiple generations, consistent with the evidence reviewed above: if a new migrant does not switch traits immediately after migrating, they may do so in subsequent generations. It also parsimoniously assumes that the same conformist bias applies to all individuals, rather than positing special learning mechanisms for migrants and non-migrants.

Conformist acculturation is implemented following Boyd and Richerson [17], by assuming that individuals adopt traits based on their frequency among *n* randomly chosen demonstrators from their sub-population, with parameter *a* increasing the probability of adoption of the most frequent trait among those *n* demonstrators. When *a* = 0, transmission is unbiased and non-conformist. When *a* = 1 there is 100% probability of adopting the most frequent trait among the demonstrators, if there is one. Boyd and Richerson [17] modeled the case when *n* = 3 and there are two traits using binomial theorem. Here I extend their formulation to *n* > 3 and more than two traits using multinomial theorem.

If *p_i_* is the frequency of trait *i* in the sub-population after migration (see Eq. 1), then the frequency of trait *i* in that sub-population after conformist acculturation, 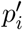, with *n* demonstrators and *s* traits, is given by:

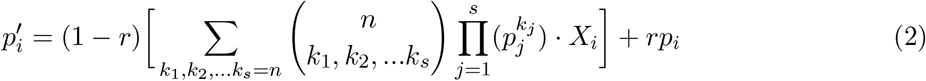

The first part of Eq. 2 gives the updated trait frequency arising from a fraction 1 − *r* of the sub-population that picks demonstrators at random. The expression in the square brackets contains the multinomial theorem and gives, for each combination of *s* traits among *n* demonstrators, the probability of that combination randomly forming and giving rise to trait *i*. *k*_1_, *k*_2_…*k_s_* represent all combinations of *s* non-negative integers from 0 to *s* such that the sum of all *k* is *n*. Conceptually, *k_j_* represents the number of demonstrators who possess trait *j*. The multinomial coefficient gives the number of ways in which demonstrators with that combination of traits can form. This is multiplied by the probability of that combination forming, with each trait frequency *p_j_* raised to the number of times that trait appears in the combination, *k_j_*, and *X_i_*, which gives the probability of adopting trait *i* for that combination of demonstrators. If *k_max_* is the maximum *k* in a combination, and *fi* is the number of traits that have *k* = *k_max_* (so when *π* = 1 there is a single most-common trait, and when *π* > 1 there are more than one joint-most-common traits), then:

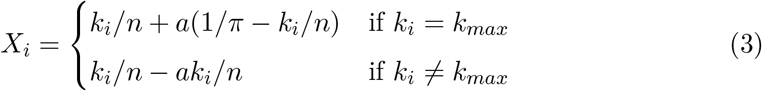

The conformity parameter *a* increases the probability with which the majority trait(s) is adopted, and decreases the probability with which minority traits are adopted.

To give a concrete example, imagine there are three traits (*s* = 3) and ten demonstrators (*n* = 10). Two demonstrators have trait 1, five have trait 2 and three have trait 3. Then, with unbiased, non-conformist transmission (*a* = 0), the probabilities of adoption of traits 1, 2 and 3 are 0.2, 0.5 and 0.3 respectively. Under weak conformity of *a* = 0.1, the probability of adopting trait 2 increases to 0.52 and the probabilities of adopting traits 1 and 3 drop to 0.19 and 0.29 respectively. Under maximum conformity (*a* = 1), the most common trait, trait 2, is adopted with probability 0.715, and traits 1 and 3 with probabilities 0.073 and 0.212 respectively. Increasing *n* also increases the effect of conformity; increasing *n* to 100 (with *a* = 1) increases the probability of adoption of the most-common trait 2 to 0.99, almost guaranteeing its adoption. Note that Equation 3 allows conformity to act on combinations where there is more than one most-common trait amongst the *n* demonstrators, by increasing all most-common traits’ frequencies equally and proportionally to *a*.

The final part of Eq. 2 deals with non-random demonstrator formation. It is reasonable to assume that, in reality, people are more likely to interact with culturally similar others. This is analogous to assortative mating. In the context of migration, this typically means that migrants will interact only with other migrants, and non-migrants with other non-migrants, at least initially when migrants and non-migrants are culturally dissimilar. In the model, I assume that a fraction *r* of demonstrators form such that only sets of culturally homogenous demonstrators come together. These form with probability equal to the frequency of the trait they all exhibit, *p_i_* (see S1 Methods for derivation). Assortation therefore leaves cultural trait frequencies unchanged, hence the term *rp_i_*.

The S1 Methods gives an example of how Eq. 2 and 3 yield trait frequencies following conformist acculturation and assortation. The model was implemented using numerical simulation tracking trait frequencies in each sub-population over time. As an independent replication of the results, the model was also implemented as an individual-based model in which individuals and their traits are explicitly simulated, which gave almost-identical results (see S1 Results).

## Results

### Migration without acculturation eliminates between-group variation

Without acculturation, migration always eliminates between-group variation, just as in Wright’s original island model and specified by Eq. 1. Fig 1 shows time series of *F_ST_* for low (*m* = 0.01), typical (*m* = 0.1) and high (*m* = 0.3) migration rates. In all cases, the grey *a* = 0 lines fall to zero. Each trait in each population converges on the mean frequency of that trait across the whole population. Because each sub-population is equally sized, and each trait starts out at unity in one of these s sub-populations, this mean frequency 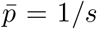 for every trait. This can occur rapidly: for *m* = 0.01, the perfect between-group structure completely disappears within 300 timesteps, while for *m* = 0.1 it disappears within 30, and for *m* = 0.3 within 10.

**Figure 1:**
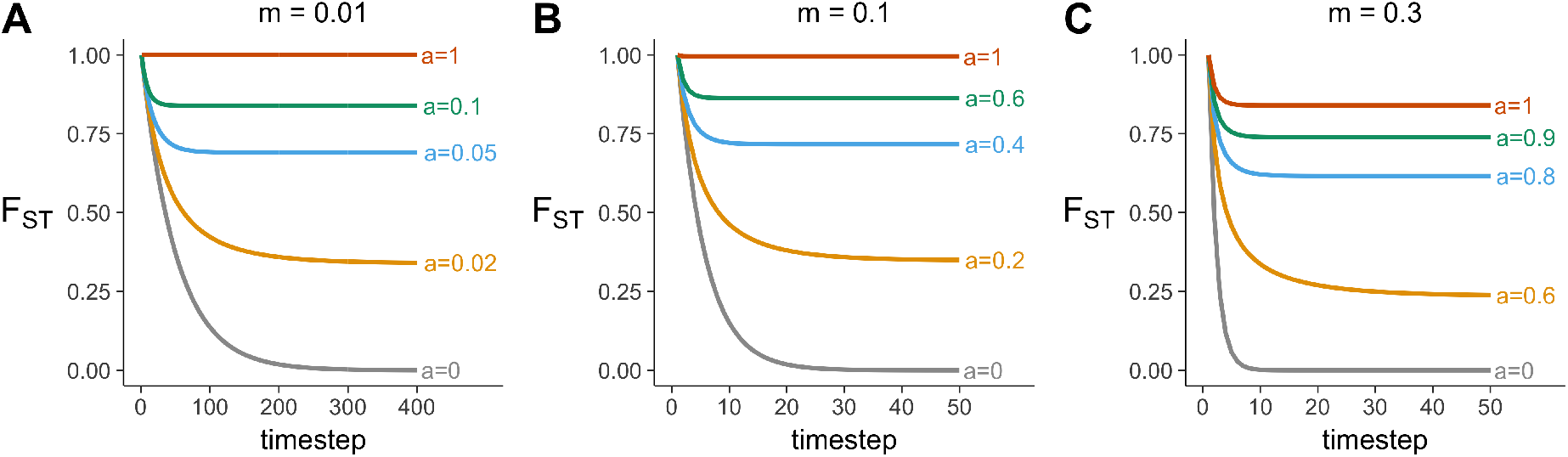
Time series showing changes in *F_ST_* over time for (A) a low migration rate *m*=0.01, (B) a typical migration rate *m*=0.1, and (C) a high migration rate *m*=0.3, at varying strengths of acculturation, *a*. Other parameters: s=5, *n*=5, *r*=0.

### Conformist acculturation can maintain between-group cultural variation

Fig 1 shows that as conformist acculturation increases in strength, between-group cultural variation is maintained (*F_ST_* > 0) at equilibrium. When migration is low or moderate then complete acculturation (*a* = 1, red lines) causes *F_ST_* to remain at 1, indicating the maintenance of complete between-group cultural variation. Values of *a* between 0 and 1 generate equilibrium values of *F_ST_* between 0 and 1, at a level at which migration and acculturation balance out. That is, the decrease in a common trait’s frequency due to migration is equal to the amount by which conformist acculturation increases the common trait’s frequency after migration. At low migration rates, even small amounts of acculturation can restore *F_ST_* to realistically high values. In Fig 1, when *m* = 0.01, then *a* = 0.1, or an extra 10% chance of adopting majority traits per timestep, maintains *F_ST_* at approximately 0.87, which is already higher than the highest *F_ST_* found by [46]. An *F_ST_* of approximately 0.35 can be maintained with just *a* = 0.02. When *m* = 0.1, then higher acculturation rates are needed to maintain between-group structure, but an *a* of just 0.2 is needed to maintain *F_ST_* at realistic levels of around 0.35. At high migration rates (*m* = 0.3), the maximum strength of conformist acculturation (*a* = 1) does not maintain complete between-group cultural variation, and higher values of *a* are needed to prevent the loss of all between-group cultural variation. Nevertheless, even with such high migration, conformist acculturation can still maintain plausible levels of between-group variation.

Fig 2 shows how the full range of *a* affects *F_ST_*, for three different migration rates and four different values of *n*. Fig 2 confirms that higher *F_ST_* can be maintained with stronger acculturation (larger *a*) and lower migration (smaller m). For high migration rates (here, *m* = 0.1 and *m* = 0.3), equilibrium *F_ST_* also increases with *n*. At low migration rates (*m* = 0.01), *n* = 3 is just as effective as larger values of *n*. This is because the migration rate is so low as to maintain homogeneity in demonstrators no matter how small the sample. When migration is high (*m* = 0.3), the minimum number of demonstrators that are required for conformity to work, *n* = 3, fails to maintain any between-group variation at any value of *a*. This shows that conformist acculturation crucially depends on *n* as well as *a*.

**Figure 2:**
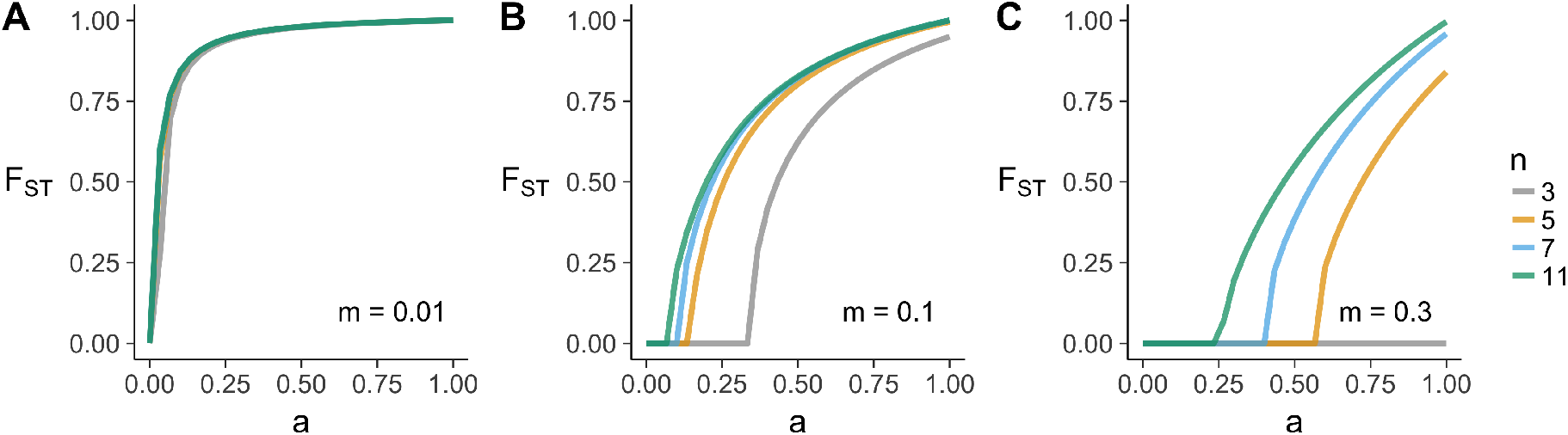
Equilibrium *F_ST_* as a function of acculturation strength, *a*, at three different migration rates, *m* (panels A-C), and different numbers of demonstrators, *n* (colored lines). Other parameters: *s*=5, *r*=0; plotted are values after 500 timesteps.

The top row of Fig 3 shows how *FST* varies across the full *m* and *a* parameter space, for two values of n. Higher values of *F_ST_* are maintained with stronger acculturation (higher *a*) and lower migration (smaller *m*). Increasing *n* from *n* = 3 to *n* = 7 increases the parameter space in which *F_ST_* is maintained above zero. This is as expected. When *n* = 3, there is little opportunity for conformity to act (only when two demonstrators exhibit one trait and one demonstrator exhibits another), while for *n* = 7 there are more opportunities (demonstrator trait ratios of 6:1, 5:2 and 4:3 all give majorities). For high migration rates of *m* > 0.5, no amount of acculturation can prevent the elimination of between-group cultural variation, although this level of migration is already unrealistically high. For plausible values of *m* around 0.1, a substantial range of *a* yields realistic *F_ST_*s of 0.1 or more, especially when there are many demonstrators.

**Figure 3:**
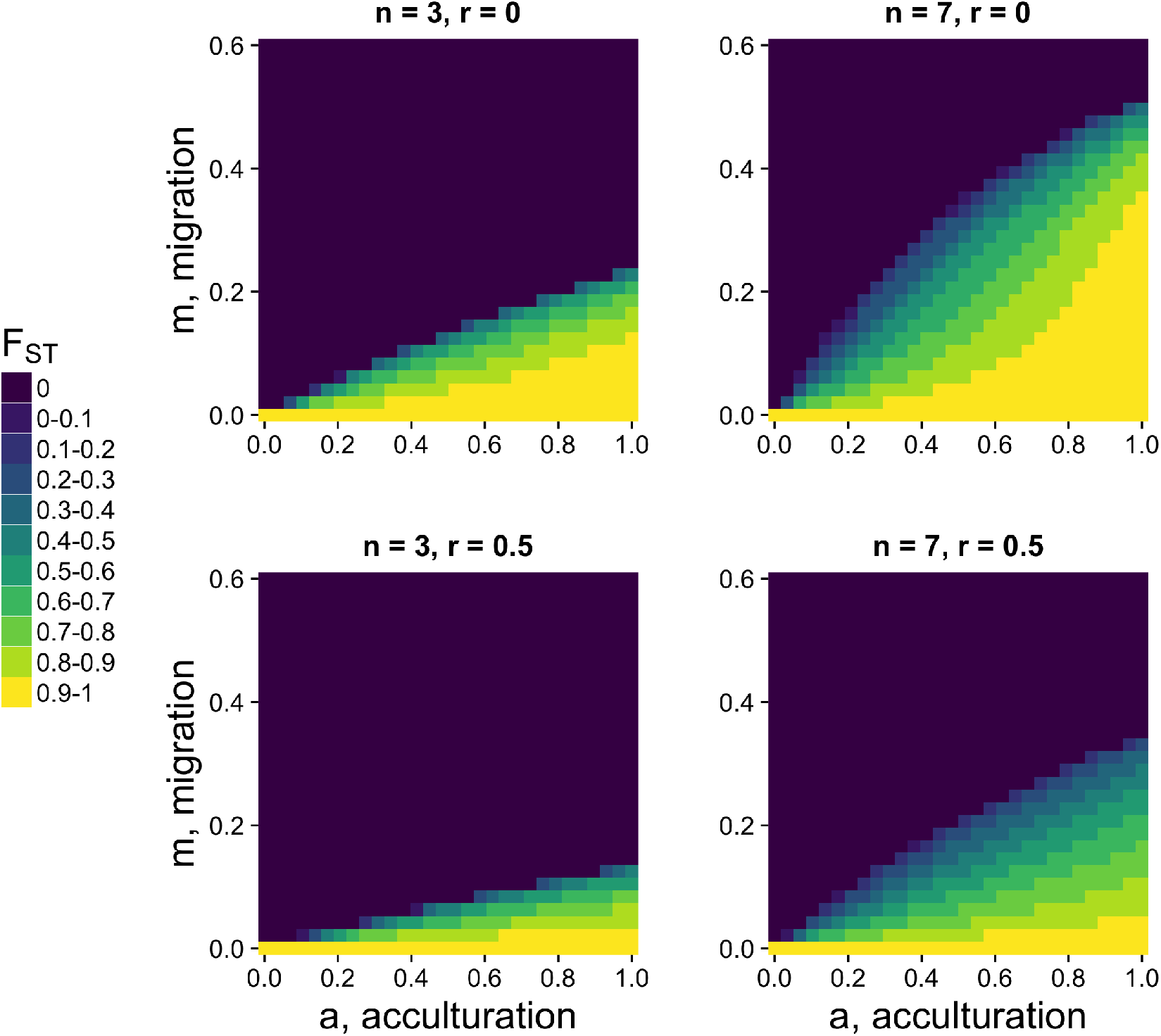
Heatmaps showing equilibrium *F_ST_* for varying acculturation rates, *a*, and migration rates, *m*, separately for different values of *n*, the number of demonstrators, and *r*, the probability of assortation. Other parameters: *s*=5; plotted are values after 500 timesteps.

### Assortation reduces the effect of acculturation

Assortation acts to restrict the parameter space within which *F_ST_* is maintained, as shown in the lower two panels in Fig 3 where *r* = 0.5. This is as expected given Eq. 2. Assortation counteracts the effect of conformity by reducing the likelihood that heterogeneous sets of demonstrators come together, which is required for conformity to operate.

### Summary of Model 1

Model 1 affords the following insights. First, even small amounts of migration can rapidly break down between-group cultural variation. This is analogous to how migration in genetic evolution can rapidly break down between-group genetic variation, and is well known from population genetics. Second, as shown in previous cultural evolution models, the breakdown of between-group cultural variation by migration can be prevented by conformist acculturation, via both the strength of this conformist bias, *a* (what might be considered a psychological factor), and the number of demonstrators from whom individuals learn, *n* (what might be considered a social or demographic factor). As both *a* and *n* increase, between-group cultural variation is more likely to be maintained. Third, at realistic and typical levels of migration - around *m* = 0.1 - even small amounts of acculturation can counteract migration and maintain plausible amounts of between-group cultural variation. At *m* = 0.1, just a small increase in the probability of adopting majority traits (*a* = 0.2) amongst a small number of demonstrators (*n* = 5) can maintain plausible amounts of between-group cultural variation. Even the highest levels of migration currently seen in the real world - around *m* = 0.3 - can still be prevented from breaking down between-group variation with moderately strong acculturation. Although there is a paucity of empirical data on the individual-level mechanism of acculturation, existing acculturation research (see Table S1 and Fig S1) suggests that acculturation is easily strong enough to meet this criterion, although there is a need for more research along the lines of [19] that directly estimates *a* across multiple migrant generations, and tests whether acculturation really is conformist. Finally, assortation acts against acculturation: when individuals only learn from culturally homogenous sets of demonstrators, then acculturation is weaker. This reflects the common intuition that more integrated migrant minorities are more likely to acculturate than marginalized minorities.

## Model 2: Migration, acculturation and cooperation

Model 1 examined the effects of migration and acculturation on neutral cultural variation. Traits were equally likely to be copied and did not affect individuals’ fitness. Migration was also random between groups. Much public concern over migration, however, centers on cooperative behavior and norms, for which neither of these assumptions are true. Think of concerns among some US citizens over Mexican immigration, and some Europeans over North African or Middle Eastern immigrants. In such cases, people are concerned that migrants coming from societies with less-cooperative norms (e.g. high crime, tax avoidance and corruption; military conflict; weak or corrupt institutions) moving to societies with more-cooperative norms (e.g. low crime, tax avoidance and corruption; strong institutions such as public health or education) will bring their less-cooperative norms with them, free-ride on the more-cooperative norms of their adopted society, and cause a break-down of cooperation in the adopted society. Migration is not random here: people are more likely to move from less-cooperative to more-cooperative societies [52,53], which will exacerbate the degradation of cooperation in the latter.

This situation again concerns acculturation. The above scenario assumes that migrants bring and retain their non-cooperative norms with them, but in reality they may acculturate to the cooperative norms of the adopted country. But how much acculturation is enough to maintain cooperative norms? Dimant et al. [54] found that migration in general has no effect on a receiving country’s corruption level, suggesting that migrants acculturate to such an extent that between-group variation in corruption is maintained. However, when examining migration only from highly corrupt countries, there was a small positive effect on the receiving country’s corruption levels, although this effect attentuated over time. This indicates that corruption can be transmitted by migrants, but acculturation may counteract this over successive generations. This is consistent with the findings of [32] (see Table S1), which found that second generation migrants in high-trust countries typically have institutional trust that is intermediate between their parents and their birth countries.

Human cooperation is also of major scientific interest, with much debate concerning how to explain the unusually high levels of non-kin-directed altruism in humans [46,55,56]. Typically, migration is seen as a force acting against cooperation because it breaks up groups of cooperators and spreads selfish free-riding behavior [55,57]. Theories of cultural group selection require stable between-group cultural variation in cooperative behavior and so need some acculturating mechanism to work against migration [46].

Model 2 therefore examines the effect of migration and acculturation on the maintenance of a cooperative cultural trait in the face of incoming migrants with non-cooperative norms. While this is an extreme case, it is useful for delineating the effect of different forces. Additional parameters in Model 2 are listed in Table 2.

**Table 2:**
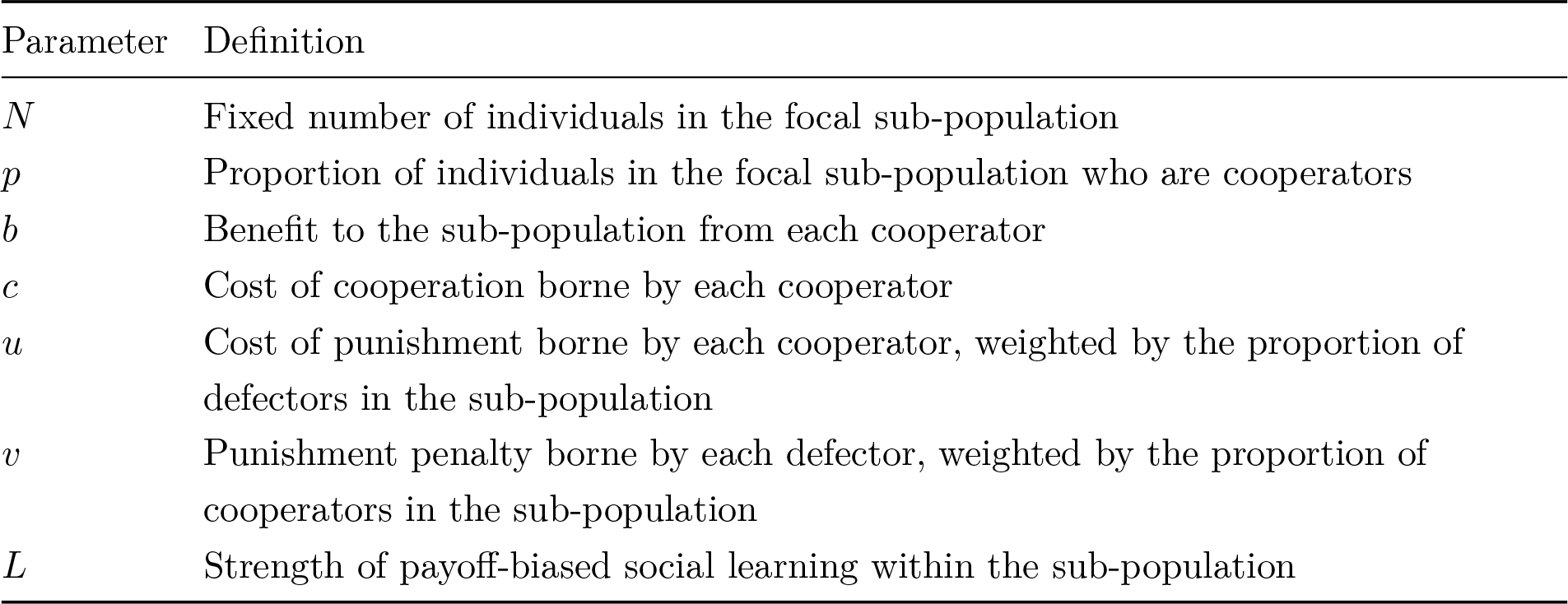
Additional parameters in Model 2

| Parameter | Definition |
| --- | --- |
| $N$ | Fixed number of individuals in the focal sub-population |
| $p$ | Proportion of individuals in the focal sub-population who are cooperators |
| $b$ | Benefit to the sub-population from each cooperator |
| $c$ | Cost of cooperation borne by each cooperator |
| $u$ | Cost of punishment borne by each cooperator, weighted by the proportion of defectors in the sub-population |
| $v$ | Punishment penalty borne by each defector, weighted by the proportion of cooperators in the sub-population |
| $L$ | Strength of payoff-biased social learning within the sub-population |

Individuals are either cooperators or defectors, and are in sub-populations of constant and equal size *N*. We are interested in the maintenance of cooperation in a sub-population where cooperation is common yet faces migrants coming from sub-populations where defection is common. Assume for simplicity a single focal sub-population initially composed entirely of cooperators (*p* = 1, where *p* is the proportion of cooperators), surrounded by a larger meta-population that supplies defecting migrants and which is so large as to have a fixed *p* = 0.

Within the focal sub-population, in each timestep each cooperator pays a cost *c* (*c* > 0) to benefit the entire sub-population by an amount *b*, where *b* > *c*. Defectors pay no cost and give no benefit. The total group benefit in the sub-population, *bNp*, is divided equally among all *N* sub-population members. Cooperators in the sub-population therefore have fitness *w_c_* = 1 + *bp* − *c* and defectors have fitness *w_d_* = 1 + *bp*, where 1 is baseline fitness.

Defectors will always have higher fitness than cooperators for *c* > 0 and always go to fixation, assuming some selective force such as payoff-biased social learning (see below) or natural selection. As soon as mutation, errors or migration introduce defectors into the cooperating group, cooperation will disappear. This is unrealistic for most human groups and makes the present model uninteresting. I therefore introduce a mechanism to maintain cooperation: coordinated altruistic (i.e. costly) punishment. Punishment is a common strategy for maintaining cooperation, and may arise via trial-and-error to create institutions [58], between-group selection [55] or other mechanisms. I am not concerned here with these processes, and assume that punishment has already evolved.

Hence, assume each cooperator pays a cost *u/N* per defector to reduce the payoff of each defector by *v/N*, where *v* > *u* [55]. There are *Np* cooperators who punish each defector, so defectors now have overall fitness of *w_d_* = 1 + *bp* − *vp*. Each cooperator punishes *N*(1 − *p*) defectors, so cooperators have fitness *w_c_* = 1 + *bp* − *c* − *u*(1 − *p*). Cooperators and defectors will have equal fitness when *w_d_* = *w_c_*, or when *p* = *p*\*, where

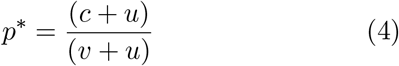

Defectors will invade a population of cooperators when *p* < *p*\*. That is, cooperation is maintained when cooperators are common enough that the punishment costs to defectors outweigh the costs to cooperators of cooperating. When *c* > *v*, cooperation is never maintained. Note that non-punishing cooperators could invade a population of punishing cooperators because the former would not pay the cost *u*. I assume that this second-order free-riding problem is already solved (e.g. by the mechanisms above) and non-punishing cooperators are not included in the model. I also assume that a sub-population entirely composed of defectors (*p* = 0) always has lower fitness than a sub-population with any cooperators (*p* > 0). See S1 Methods for details.

As in Model 1, the first timestep event is migration determined by migration rate m, but this is now payoff-biased. Individuals move to sub-populations with higher mean fitness than their own at a rate proportional to the mean fitness difference between their own sub-population and the meta-population [52]. By assumption, in Model 2 this will always involve defectors coming into the sub-population and replacing cooperators (who are assumed to die or migrate away) at a rate proportional to *W* − 1, where *W* is the mean fitness of the focal sub-population and 1 is the mean fitness of the meta-population where *p* = 0. The frequency of cooperators in the sub-population after migration, *p′*, is therefore

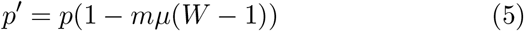

The constant *μ* ensures that *mμ*(*W* − 1) never exceeds *m*, so that *m* is always the maximum migration rate. Eq. 5 resembles Eq. 1, but with *m* now a function of relative sub-population fitness, and 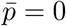.

Following migration there is acculturation, identical to Model 1. With probability *a*, each individual adopts the most common strategy (cooperate or defect) among *n* demonstrators in their subpopulation according to Eq. 2 (with *s* = 2, given two traits, cooperate and defect). This occurs after all migration has finished.

Finally there is payoff-biased social learning within each sub-population. With probability *L*, individuals switch strategies in proportion to the fitness payoff difference within their sub-population between the alternative strategy and their current strategy. If *p′* is the frequency of cooperators after migration and conformist acculturation, then the frequency after payoff-biased social learning, *p″*, is given by:

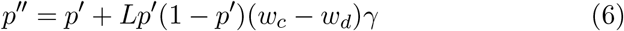

where *γ* is a constant that scales *L* according to the maximum possible fitness difference. Payoff-biased social learning creates a selective force within the sub-population favoring whichever strategy gives the highest payoff, which in turn depends on Eq. 4.

Model 2 comprises cycles of Eq.s 5, 2 and 6 (payoff-biased migration, conformist acculturation and payoff-biased social learning). As we are interested in the maintenance of cooperation, we track the proportion of cooperators *p* over time in the focal sub-population which initially comprises all cooperators.

## Results

### Payoff-biased migration alone eliminates cooperation

In the absence of acculturation (*a* = 0) and payoff-biased social learning (*L* = 0), payoff-biased migration (*m* > 0) causes defectors to flow from the all-defect meta-population into the initially all-cooperate sub-population to eliminate cooperation entirely (Fig 4A). Because the strength of payoff-biased migration is a function of the mean population fitness relative to the mean fitness of the metapopulation, the rate of decline is initially fast due to the high initial mean fitness of the cooperative sub-population, and slows as cooperators leave and mean fitness drops.

**Figure 4:**
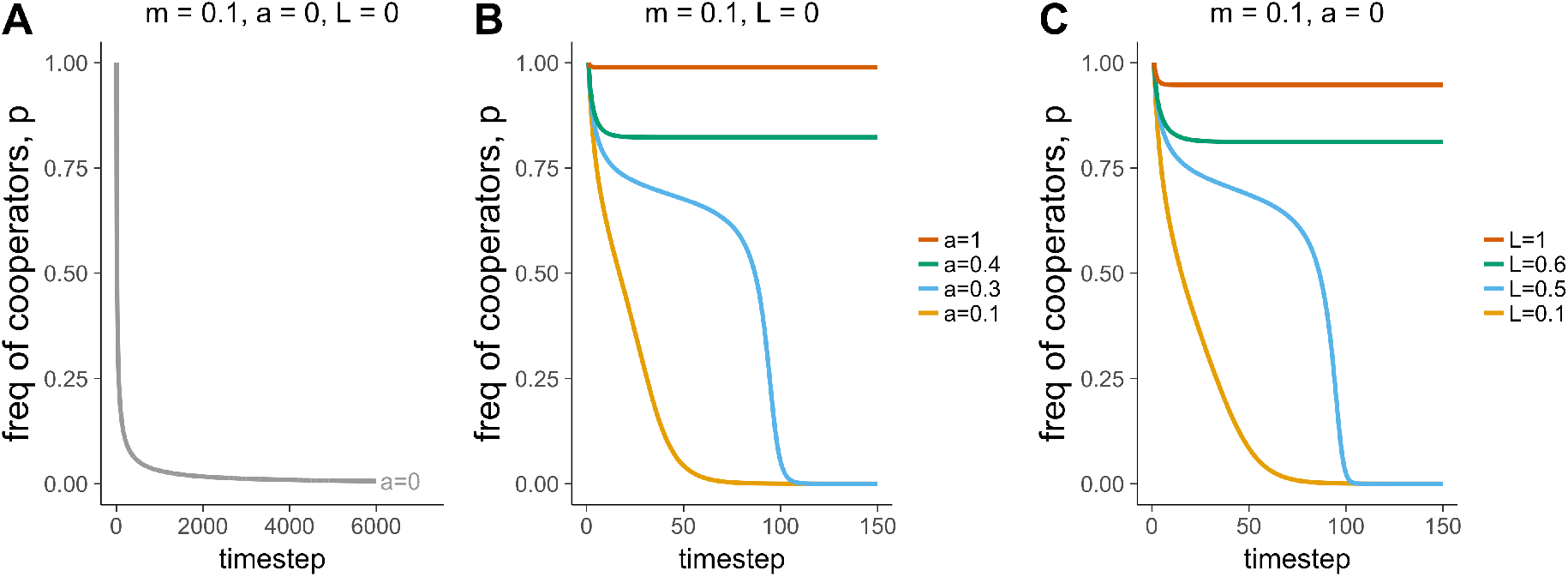
Time series showing changes in *p* over time in the face of payoff-biased migration (*m*=0.1), (A) in the absence of acculturation (*a*=0) and payoff-biased social learning (*L*=0); (B) at varying strengths of acculturation, *a*, and (C) at varying strengths of payoff-biased social learning, *L*. Other parameters: *n*=5, *r*=0, *b*=1, *c*=0.2, *u*=0.1, *v*=0.5.

**Figure 5:**
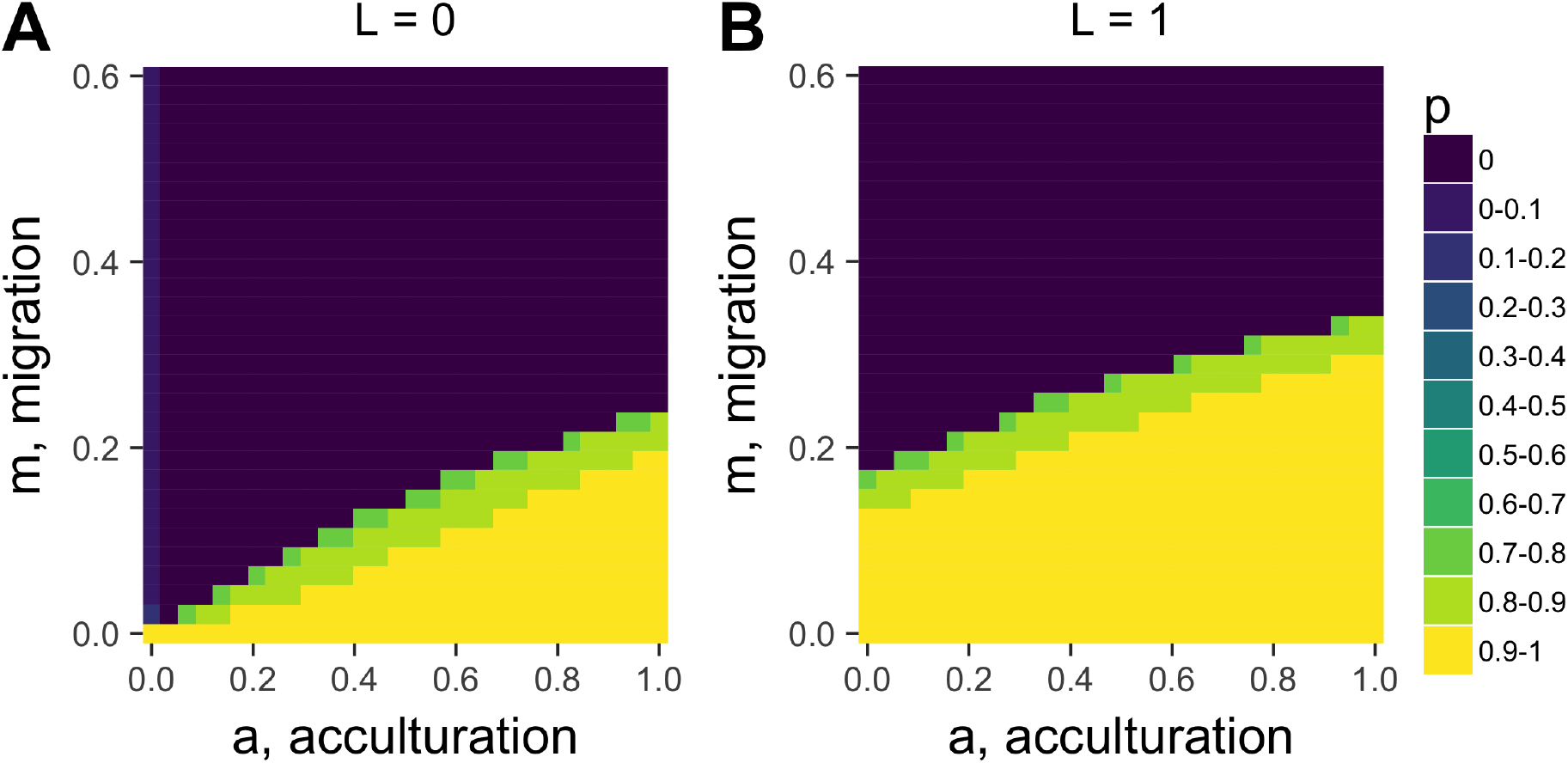
Heatmaps showing equilibrium *p* for varying acculturation rates, *a*, and migration rates, *m*, separately for different values of *L*, the strength of payoff-biased social learning. Other parameters: *r*=0, *n*=5, *b*=1, *c*=0.2, *u*=0.1, *v*=0.5; plotted are values after 1000 timesteps.

### Conformist acculturation can maintain cooperation

As in Model 1, when conformist acculturation is sufficiently strong (i.e. *a* and *n* are sufficiently large), then the decline in cooperation is halted and cooperation is maintained at a point where acculturation and migration balance out (Fig 4B). This can also be seen in Fig 5A, which shows a similar relationship between *a* and *m* as in Model 1: cooperation is most likely to be maintained when *a* is high, and *m* is low.

Two points are worth noting. First, when acculturation is not strong enough to maintain cooperation, it actually speeds up the decline. Compare the several thousand timesteps it takes for cooperation to drop to approximately *p* = 0 in Fig 4A for *a* = 0 to the 100 timesteps it takes to reach *p* = 0 in Fig 4B for *a* = 0.1. Conformity favors the majority trait, which when *p* < 0.5 is defection, speeding up the convergence on *p* = 0.

Second, unlike in Model 1, we see an interesting dynamic at values of *a* that are not strong enough to maintain cooperation (e.g. *a* = 0.3 in Fig 4B). An initial rapid decline in cooperation when *p* = 1 slows as *p* declines, then increases again. This can be understood in terms of the relative strengths of payoff-biased migration and conformist acculturation. Payoff-biased migration is strongest at *p* =1 and weakens as it approaches its stable equilibrium at *p* = 0. Conformist acculturation has an unstable equilibrium at *p* = 0.5 where the two traits are equal in frequency, and increases in strength as the frequency approaches the two stable equilibria at *p* = 0 and *p* =1. In Fig 4B when *a* = 0.3, the initial rapid decline is due to strong payoff-biased migration near *p* =1. As *p* decreases, payoff-biased migration weakens and conformist acculturation slows the decline. As we approach *p* = 0.5 conformity weakens, allowing payoff-biased migration to take over and increase the rate of decline. When *p* drops below 0.5, conformity starts to work with payoff-biased migration to increase the rate of decline further.

This dynamic means that there are fewer values of *a* in Model 2 at which *p* is maintained at intermediate values, compared to Model 1. This can be seen in Fig 5A: final frequencies are typically either *p* = 0 (dark blue) or *p* > 0.8 (yellow / light green), with little in between. Contrast this to Model 1 in Fig 3, which has more intermediate frequencies.

### Payoff-biased social learning acts like an acculturating mechanism

Fig 4C shows a strikingly similar dynamic for payoff-biased social learning (*L*) as for conformist acculturation. When *L* is large, cooperation is maintained. When *L* is small, cooperation declines to zero, with a similar-shaped curve for intermediate values of *L*. Payoff-biased social learning therefore acts as an acculturating mechanism, like conformity. It does this because cooperation is favored when common, and disfavored when less common, similar to conformity (see S1 Methods). Fig 4B shows how *L* increases the parameter space in which *a* maintains cooperation in the face of migration, showing how they work in the same manner.

**Figure 6:**
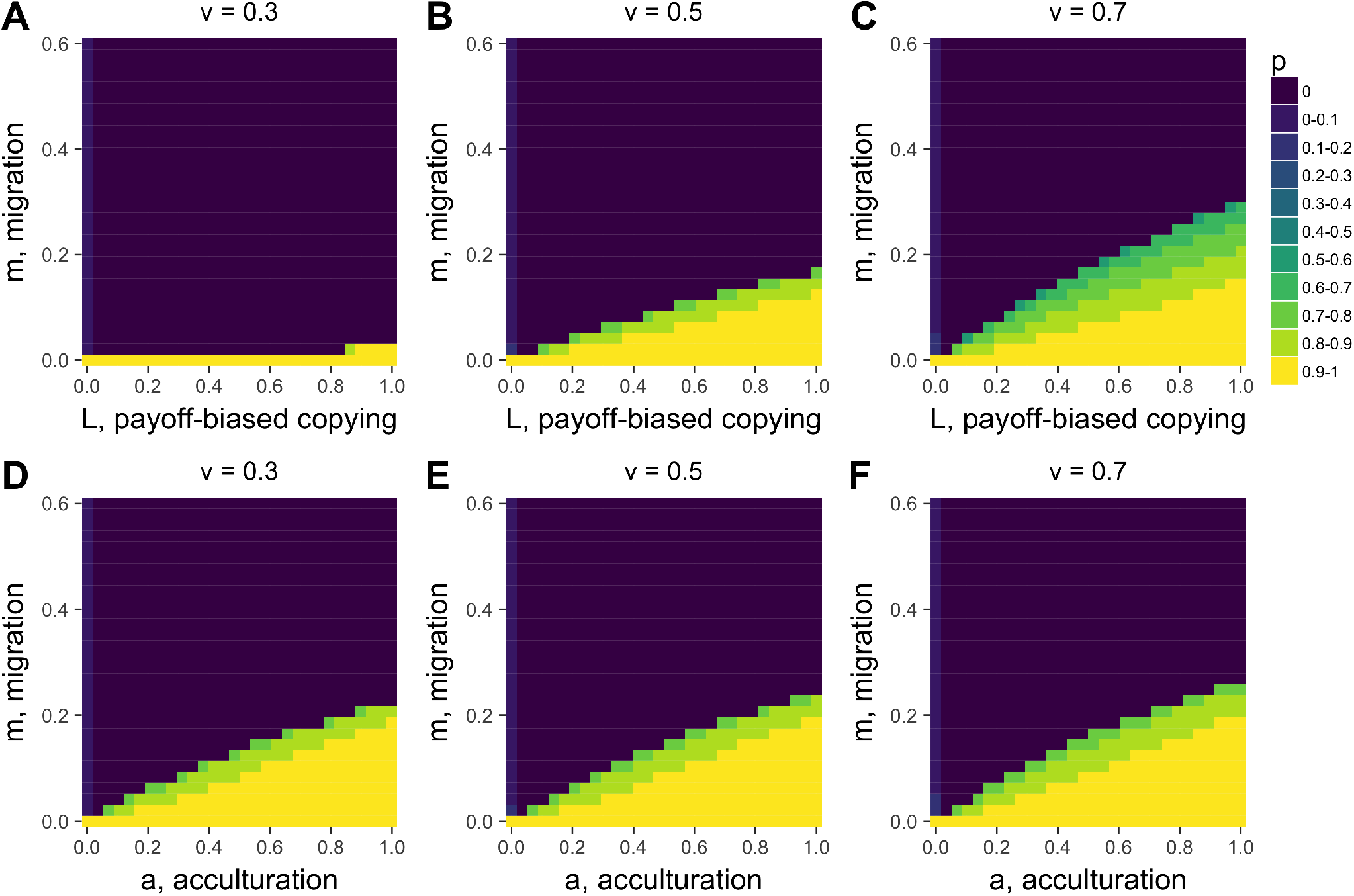
The heatmaps in A-C show how different values of *v*, the punishment costs to defectors, strongly affect the relationship between the strength of payoff-biased within-group social learning, *L*, and migration rate, *m* in determining equilibrium cooperation level p. Heatmaps D-E show, in contrast, that *v* has little effect on the relationship between *m* and acculturation, *a*. Other parameters: *r*=0, *b*=1, *c*=0.2, *u*=0.1; in A-C, *a*=0 and in D-E, *L*=0. Plotted are values after 1000 timesteps. Overall, this comparison shows that the strength of *L*, unlike *a*, depends on the strength of the sanctioning institution that governs the strength of punishment.

Whereas conformist acculturation has an unstable equilibrium at *p* = 0.5 (given that there are two traits), payoff-biased social learning has an unstable equilibrium at *p*\*. In Fig 4C, *p*\* = 0.5, so there is little difference to Fig 4B. However, at different values of *p*\*, there will be different dynamics (see S1 Methods). Fig 6 (A-C) shows how changing the basin of attraction for cooperation via the punishment penalty for defectors, *v*, and therefore *p*\* (via Eq. 4), alters the effect of *L*. When *v* is small, *p*\* is large, cooperation has a smaller basin of attraction, and *L* mostly favors defectors. When *v* is large, *p*\* is small, cooperation has a larger basin of attraction, and *L* largely favors cooperators. Conformist acculturation, by contrast, does not vary with *v* or any other fitness parameter, as it solely works on the basis of trait frequency (Fig 6D-F).

### Summary of Model 2

Like Model 1, Model 2 shows that conformist acculturation works to maintain cultural traditions in the face of migration. Rather than neutral cultural variation, here the tradition involves cooperative behavior that is lost due to payoff-biased migration bringing non-cooperators from low-payoff to high-payoff sub-populations. The evidence reviewed above and in Table S1 includes cases of acculturation of cooperative behavior, with migrants from low-trust or high-corruption countries acculturating to the high-trust or low-corruption norms of their adopted country within a few generations [32,54]. More empirical work is required to accurately specify acculturation rates across a broader range of societies and norms, and test whether it is conformist. However, we can apply the same insight here as from Model 1: surprisingly little acculturation is needed to maintain cooperation in the face of realistic levels of migration. Fig 4B and 5A, for example, show that for *m* = 0.1, an *a* of around 0.4 (with *n* = 5) is sufficient to maintain cooperation. On the other hand, where migration *is* strong enough to allow non-cooperating migrants to increase in frequency, conformity speeds up the loss of cooperation when cooperation becomes a minority trait. There are also fewer intermediate cases: cooperation is typically either maintained at a high level, or lost completely.

Finally, within-group payoff-biased social learning plus punishment acts like an acculturating process, also maintaining cooperation in the face of migration. This is because cooperation is frequency dependent, with cooperators having high fitness when common due to punishment. Unlike conformity, the acculturating power of payoff-biased social learning depends on the fitness functions and the strength of punishment. The real-world correlate of the latter might be the strength of sanctioning institutions. When punishment of defectors is more effective via higher *v* (stronger sanctioning institutions), then payoff-biased social learning more effectively prevents migration from eliminating cooperation.

## Discussion

Public debates concerning migration often proceed with little empirical basis. Academic research into migration is voluminous but seldom seeks to formally link individual-level learning, interaction and acculturation to population-level patterns of cultural diversity and change. I have attempted here to formalise the processes that contribute to the maintenance and loss of cultural diversity in the face of migration, drawing on and extending previous models of cultural evolution. Migration itself (*m*) breaks down between-group cultural variation and homogenises behavior. Acculturation, here assumed to be conformist, prevents this breakdown. This acculturation effect increases with both the strength of the conformist learning bias (*a*) and the number of individuals from whom one learns (*n*), but weakens when individuals interact assortatively with culturally similar others (*r*). The learning bias strength might be considered a psychological factor, while the number of demonstrators and assortation might be considered social or demographic factors. Model 1 found that surprisingly little acculturation is needed to maintain between-group variation in functionally neutral cultural traits given plausible rates of migration. Model 2 found similar results for a cooperative cultural trait, which is additionally subject to fitness costs and benefits. Even though payoff-biased migration acts to reduce cooperation, conformist acculturation and/or payoff-biased social learning (the latter working together with sanctioning institutions that punish non-cooperators) can preserve cooperation.

How do these insights relate to the empirical data concerning migration and acculturation? The evidence regarding the strength of acculturation reviewed above (see also Table S1 and Fig S1) suggests that migrants often shift around 50% towards the values of their adopted society every generation, although this varies across traits and societies. In the context of the models, we can tentatively conclude that these acculturation rates are easily strong enough to maintain cultural traditions in the face of migration. The models suggest that values of *a* much less than 0.5 can preserve variation for realistic values of *m*. If this is the case, then common fears that migration will inevitably destroy existing cultural traditions may be exaggerated or unfounded. Yet this is only a very loose comparison. There is little quantitative work that tries to measure acculturation rates in a way comparable to acculturation as modelled here. We do not know how migrants integrate social information from multiple sources, whether they do this in a conformist manner, the size and diversity of their social networks, or how this changes with successive generations who experience different social environments compared to their parents. Perhaps demonstrator-based biases, where people preferentially learn from certain sources such as parents, teachers or celebrities [17], are more appropriate learning mechanisms. We also do not know whether acculturation of cooperative norms is driven by conformist or payoff-biased social learning, both of which have similar acculturating effects but for different reasons.

While the models here might counter extreme conservative claims that *any* level of migration is detrimental for the maintenance of cultural traditions, they also count against extreme liberal claims that migration can never be too high. For very high rates of migration (e.g. *m* > 0.5) then between-group cultural variation is typically eroded completely. While such levels exceed modern-day migration rates, such a situation might resemble past colonisation events. The colonisation of the New World, for example, led to the elimination or attenuation of much pre-contact cultural variation and replacement with European cultural values such as religious beliefs. Further models might integrate such power-based dynamics into the framework developed in the present paper.

The models presented here necessarily comprise many simplifying assumptions and omissions that could be addressed in future models. The present models failed to incorporate any benefits of migration, such as when migrants bring beneficial new skills and knowledge into a society and recombine them with existing skills and knowledge. The assumption of Model 2 that migrants always possess non-cooperative norms is obviously an unrealistic extreme case. Future models might examine the two-way exchange of cooperative behavior between multiple sub-populations, and the consequences for between-group competition by allowing *N* to vary as a function of mean sub-population fitness [52]. The present models only allowed individuals to hold one trait at a time. Future models might examine the simultaneous acculturation of different traits at different rates, as found empirically [19], and the consequences for multi-dimensional cultural diversity. Finally, the traits here are discrete whereas many cross-cultural measures are continuous, which may gradually shift over time due to averaging across different demonstrators [17].

Note that acculturation as considered in the models in the current study is considered in the restricted sense of the extent to which migrants become more similar to the adopted population, or retain their heritage values, on a single quantitative dimension. This is only one possible acculturation dynamic. In Berry’s [15] classification of acculturation styles, this is the *assimilation* vs *separation* dimension, the former involving shifts in migrants towards host values, the latter involving retention of heritage values. Berry [15] also highlights the possibility of *integration*, which entails maintaining heritage traits *and* adopting host society traits and possibly merging the two; and *marginalization*, which entails the rejection of both heritage and host society traits. Similarly, political scientists categorize modern nation state policies on migration as *exclusionary*, *assimilationist* or *multicultural* [53]. The *integration* and *multicultural* dynamics in these schemes serve to highlight how migrants can contribute to and shape their adopted country’s culture. Migration is a complex phenomenon, and the empirical literature reviewed here, and the models in this study, represent an attempt to understand one particular simplified aspect of this complex reality.

One recent paper presents similar models of acculturation and migration [20], drawing on Berry’s [15] classification of acculturation styles. In that model, acculturation was implemented as a probability of switching to another individual’s cultural trait given the probability of those two individuals interacting plus the individuals’ relative cultural conservatism. This is an alternative to the conformity modelled here; one key difference is that in [20] residents and immigrants have different levels of conservatism, whereas here residents and migrants have identical conformist biases. That study also only modelled a single focal population, rather than multiple sub-populations and traits as in Model 1 here (although similar to my Model 2). The focus there was on when “multicultural” societies emerge; here it is on when between-group diversity persists. Nevertheless, broadly similar findings can be observed: there, resident traits disappear when migration is high, acculturation is weak (i.e. residents and immigrants are similarly conservative) and assortation is high, which is also when between-group diversity disappears in my models. Another recent model [59] examined the maintenance of between-group cultural variation when groups differ in power, finding that variation is maintained when inter-group borders allow members of the less powerful minority group to cross but not members of the more powerful majority, or when the minority inhabit a region that the majority does not visit. In [59], however, inter-group interactions do not take the form of permanent migration, and again only two groups and two traits are considered. I anticipate a class of models such as these with different assumptions against which empirical data can be compared.

The focus here is on humans, but the same principles likely also apply to non-human culture. Between-group cultural variation has been demonstrated in numerous species [60], and conformity in at least one [42]. Field experiments have demonstrated how migrant vervet monkeys switch to locally common food choices [61], an instance of non-human acculturation. Integrating human migration research with this non-human research, which often has far better individual-level behavioral data [42], will be beneficial to each.

If conformity is the mechanism underlying migrant acculturation, it is instructive to consider how conformity itself might vary between individuals and societies. Evidence suggests that social learning biases such as conformity vary not only individually but also culturally [43]. Classic social psychology research suggests that East Asian societies are more conformist than Western societies [62] (albeit using a different sense of conformity to that implemented here). If that is indeed the case, then perhaps East Asian migrants with a conformist heritage would more effectively acculturate into Western society than vice versa. This might suggest that the acculturation literature, which is biased towards Asian American samples (see Table S1), is not representative of acculturation in general. On the other hand, perhaps conformity within East Asian societies does not translate to conformity amongst East Asian migrants living in Western societies. Or, perhaps conformity is itself a trait that is subject to acculturation, and second generation East Asian migrants become less conformist by conforming to Western non-conformity. This ‘social learning of social learning strategies’ deserves further modelling (e.g. [63]) and empirical study [43].

Migration is one of the fundamental drivers of genetic evolutionary change, along with selection, mutation and drift. Migration is surely also a fundamental factor driving cultural evolution, yet works differently due to the often-rapid acculturation of migrants to local behavior. A major scientific and applied challenge for cultural evolution research is to explore the role of migration and acculturation in shaping human cultural diversity both past and present. The models here provide a step towards this goal, but also highlight major gaps in our empirical knowledge regarding how acculturation operates at the individual level, which is crucial for anticipating the population-level consequences of migration on cultural change and variation.

## Acknowledgements

I am grateful to Ruan Kilroy for locating some of the studies in Table S1, and devising the measure of acculturation used in Fig S1.

## Supporting Information

**Table S1:** Empirical patterns of acculturation. For most measures, migrants are intermediate between heritage and host values (where available), and/or 2nd generation migrants are closer to host values than 1st generation migrants. Either of these indicate acculturation.

| Trait | Host country | 2nd generation | 1st generation | Heritage country | Reference |
| --- | --- | --- | --- | --- | --- |
| Collectivism | UK: 5.45 (sd=0.77, n=99) | 5.93 (sd=0.66, n=79) | 6.29 (sd=0.57, n=108) | Bangladesh: NA | [1] |
| Dispositional attribution | UK: 5.27 (sd=0.88, n=99) | 5.14 (sd=0.79, n=79) | 4.94 (sd=1.01, n=108) | Bangladesh: NA | [1] |
| Situational attribution | UK: 4.38 (sd=1.01, n=99) | 4.87 (sd=1.04, n=79) | 5.16 (sd=1.08, n=108) | Bangladesh: NA | [1] |
| Social closeness | UK: 4.18 (sd=1.79, n=99) | 4.84 (sd=1.44, n=79) | 4.93 (sd=1.87, n=108) | Bangladesh: NA | [1] |
| Religiosity | UK: 1.86 (sd=1.21, n=99) | 4.10 (sd=1.44, n=79) | 4.83 (sd=1.34, n=108) | Bangladesh: NA | [1] |
| Family contact | UK: 3.09 (sd=2.44, n=99) | 7.11 (sd=4.83, n=79) | 7.33 (sd=5.23, n=108) | Bangladesh: NA | [1] |
| Self-serving bias | Canada: 0.36 (sd=0.65, n=98) | 0.22 (sd=0.79, n=111) | NA | Japan: -0.67 (sd=1.05, n=222) | [2] |
| Self-serving bias | USA: 0.94 (n=35) | 0.69 (n=28) | NA | Japan: 0.56 (n=23) | [3] |
| Friend-serving bias | USA: 0.84 (n=35) | 0.68 (n=28) | NA | Japan: 1.01 (n=23) | [3] |
| Self-esteem | UK: 16.33 (sd=4.19, n=381) | 16.87 (sd=3.99, n=562) | NA | Hong Kong: 13.57 (sd=4.45, n=360) | [4] |
| Self-esteem (self-report) | USA: 5.36 (sd=0.93, n=166) | 4.86 (sd=1.07, n=195) | NA | China: 4.72 (sd=0.96, n=153) | [5] |
| Self-esteem (spontaneous) | Canada: 3.73 (sd=4.44, n=110) | 2.99 (sd=3.71, n=100) | NA | China: 1.71 (sd=2.02, n=95) | [5] |
| Dialectic self-perception | USA: 3.61 (sd=0.83, n=115) | 3.89 (sd=0.65, n=129) | NA | China: 3.98 (sd=0.69, n=153) | [5] |
| Actual-ideal self discrepancy | Canada: 1.20 (sd=0.49, n=90) | 1.25 (sd=0.50, n=151) | NA | Japan: 1.49 (sd=0.57, n=161) | [6] |
| Self-serving bias | Canada: 0.27 (sd=0.66, n=90) | -0.03 (sd=0.68, n=151) | NA | Japan: -0.18 (sd=0.76, n=161) | [6] |
| Life satisfaction | USA: 5.12 (sd=1.18, n=170) | 4.21 (sd=1.34, n=149) | NA | China: 3.38 (sd=1.17, n=141) | [7] |
| Self-esteem | USA: 5.77<br>(sd=0.94, n=170) | 5.09 (sd=1.18, n=149) | NA | China: 4.93<br>(sd=1.13, n=141) | [7] |
| Relationship quality | USA: 4.01<br>(sd=0.70, n=170) | 3.70 (sd=0.72, n=149) | NA | China: 3.31<br>(sd=0.86, n=141) | [7] |
| Positive affect | USA: 3.60<br>(sd=0.58, n=170) | 3.34 (sd=0.57, n=149) | NA | China: 3.07<br>(sd=0.60, n=141) | [7] |
| Negative affect | USA: 2.10<br>(sd=0.60, n=170) | 2.36 (sd=0.71, n=149) | NA | China: 2.54<br>(sd=0.66, n=141) | [7] |
| Emotional expressiveness | USA: 4.89<br>(sd=0.91, n=170) | 4.46 (sd=0.88, n=149) | NA | China: 3.95<br>(sd=0.97, n=141) | [7] |
| Emotional differentiation | USA: 4.56<br>(sd=1.06, n=170) | 4.14 (sd=0.99, n=149) | NA | China: 3.78<br>(sd=1.32, n=141) | [7] |
| Optimism | USA: 4.06 (n=257) | 3.08 (n=44) | NA | China: 2.46<br>(n=312) | [[8]] |
| Trust | Denmark: 6.80<br>(n=2616) | 6.10 (n=35) | 5.86 (n=73) | Outside Western Europe: NA | [9] |
| Trust | Norway: 6.61<br>(n=3088) | 6.31 (n=17) | 6.07 (n=73) | Outside Western Europe: NA | [9] |
| Trust | Sweden: 6.29<br>(n=3119) | 6.01 (n=63) | 5.43 (n=204) | Outside Western Europe: NA | [9] |
| Trust | Switzerland: 5.99<br>(n=2686) | 5.73 (n=82) | 5.54 (n=258) | Outside Western Europe: NA | [9] |
| Trust | Netherlands: 5.83<br>(n=3167) | 5.58 (n=126) | 5.45 (n=244) | Outside Western Europe: NA | [9] |
| Trust | Austria: 5.48<br>(n=3783) | 5.10 (n=221) | 4.98 (n=192) | Outside Western Europe: NA | [9] |
| Trust | Belgium: 5.11<br>(n=2909) | 4.46 (n=99) | 4.71 (n=125) | Outside<br>Western<br>Europe:<br>NA | [9] |
| Trust | France: 4.94<br>(n=3026) | 4.75 (n=194) | 4.72 (n=189) | Outside<br>Western<br>Europe:<br>NA | [9] |
| Trust | Greece: 3.49<br>(n=1975) | 3.69 (n=169) | 4.27 (n=175) | Outside<br>Western<br>Europe:<br>NA | [9] |

**Figure S1:**
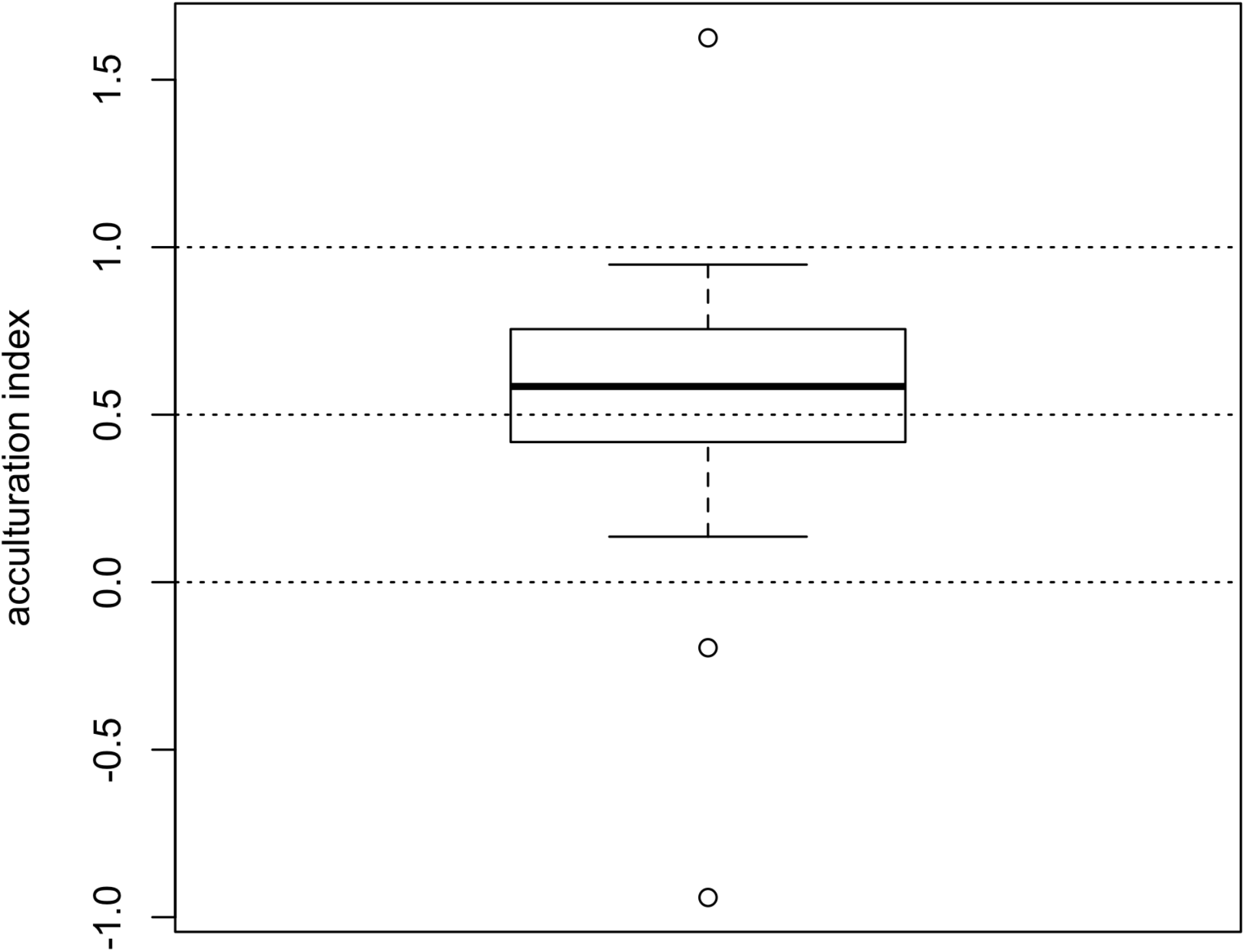
Boxplot of acculturation rates. The values in Table S1 can be used to create a crude acculturation index, representing the extent of the shift in 2nd generation migrants from their 1st generation parents or heritage countries, to their host country. This is calculated as (2nd gen - host) / (1st gen or heritage - host). A value of zero indicates no acculturation, and the 2nd generation is identical to the 1st generation or heritage population. A value of one indicates complete acculturation, and the 2nd generation are identical to the host population. Note that values can fall outside 0-1 if the 2nd generation are more extreme than the 1st generation / heritage or host populations. Dotted lines show 0 (no acculturation), 1 (complete acculturation) and 0.5 (a shift 50% towards host values). This index has mean 0.54 and median 0.58, indicating a shift in 2nd generation migrants just over half of the way from their parental generation to the host country. This measure is crude given the non-systematic selection of studies for inclusion in Table S1, and the lack of incorporation of measures of uncertainty and potentially different underlying disrtibutions for each Table S1 value used to create the index.

## S1 Methods

### Calculation of *F_ST_*

Wright’s *F_ST_* is the proportion of total population variation that occurs between sub-populations rather than within. In our models we have a population divided into *s* equally-sized sub-populations and s different traits. To calculate *F_ST_* we first calculate the total variance, i.e. the probability that two randomly chosen individuals from the entire population have the same trait, ignoring subpopulation structure. If *X_i,j_* is the frequency of trait *i* in sub-population *j*, and 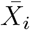 is the mean frequency of trait *i* across all sub-populations, then the total variance, *var_total_*, is given by

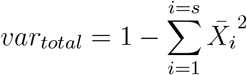

We then calculate the within-group variance for each sub-population, i.e. the probability that two randomly chosen individuals from that sub-population have the same trait. If *var_j_* is the variance in sub-population *j*, given by

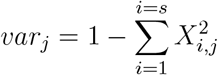

then the overall within-population variance, *var_within_*, is the mean of these variances:

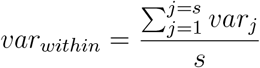

*F_ST_* is then given by

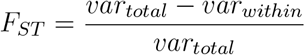

### Conformity for more than two traits and more than three demonstrators

Boyd and Richerson [10] provide a model for the change in trait frequency under the assumption of conformist transmission, such that traits that are more frequent are more likely to be adopted relative to unbiased transmission. Their basic model (on p.208) assumes two traits (*c* and *d*) and three demonstrators (they call these ‘models’ but this is confusing, so I use ‘demonstrators’). They use the binomial theorem to calculate the probability that different sets of three demonstrators will meet at random and pass on their traits. A parameter D, equivalent to *a* in the current models so henceforth labelled *a*, specifies the increased probability of adopting the majority trait (held by 2/3 of the demonstrators) and decreased probability of adopting the minority trait (held by 1/3 of the demonstrators), when demonstrators possessed different traits. When *a* = 1 there is maximum conformity, when *a* = 0 there is no conformity and transmission is unbiased. For two traits and three demonstrators, the frequency *p′* of a trait after conformity is given by

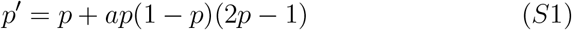

where *p* is the frequency of the trait before conformity. This however only applies when there are three randomly chosen demonstrators and two traits. The effect of increasing *a* can be seen in Figure S2. Increasing *a* from 0.5 to 1 increases the speed with which the initially more common trait goes to fixation.

**Figure S2:**
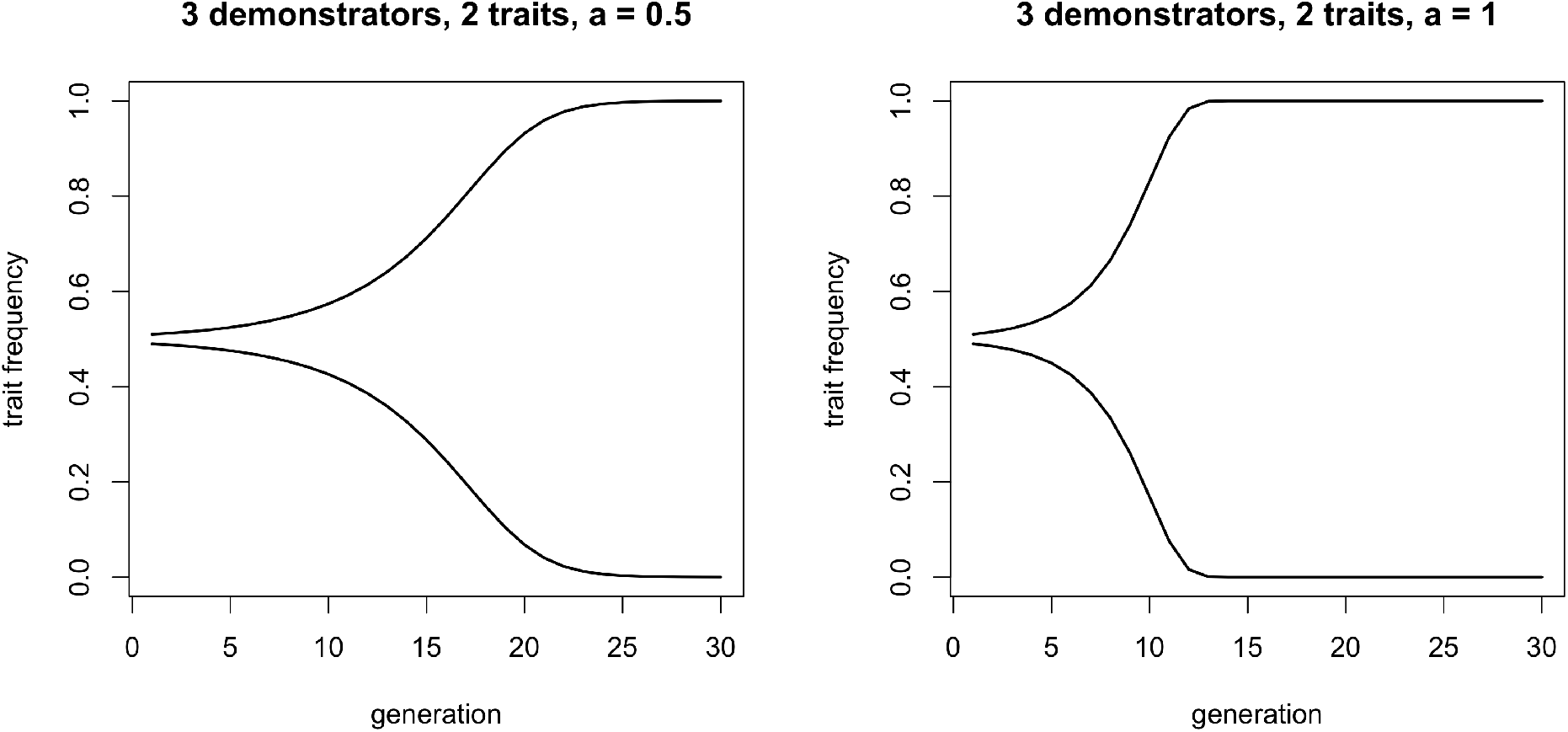
The effect of conformity on trait frequencies, for two traits and three demonstrators. The trait that is initially more common (initial frequency = 0.51) than the other trait (initial frequency = 0.49) goes to fixation. This occurs faster when the strength of conformity, *a*, is larger.

Boyd and Richerson [10] extended their model to include more than three demonstrators, but their formulation (p.213) failed to specify a conformity function. Efferson et al. [11] provided a clearer model of conformity with more than three demonstrators, also using the binomial theorem, showing that the effect of conformity increases with the number of demonstrators. However they did not extend to more than two traits. Nakahashi et al. [12] devised a model extending conformity to more than two traits, but used a different formulation to the binomial model. Their model used a parameter also labelled *a*: when their *a* = 1 there is no conformity, when their *a* > 1 there is conformity, and when their *a* = ∞ there is maximum conformity. However, this model has an unclear individual level interpretation. Boyd and Richerson’s [10] conformity parameter has a clear meaning: when their parameter equals 1, then individuals faced with majority and minority traits always pick the majority trait. It is unclear, however, what Nakahashi et al’s [12] *a* = ∞ means in this context, particularly when one wants to generate empirically testable predictions regarding acculturation strengths and compare to an individual-based model.

Here I use multinomial theorem to extend Boyd and Richerson’s [10] model to more than three demonstrators and more than two traits, and with an interpretable conformity parameter.

If *p_i_* is the frequency of a trait in the population, then to obtain the frequency of that trait in the next generation, 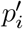, after conformist transmission with *n* randomly forming demonstrators and s traits, we can use the multinomial theorem:

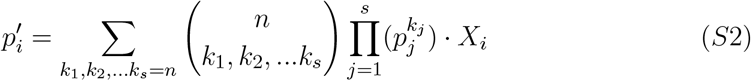

where *k*_1_, *k*_2_…*k_s_* represent all combinations of *s* non-negative integers such that the sum of all *k* values is *n*. For example, when *s* = 2 and *n* = 3, then there are two *k* values, *k*_1_ and *k*_2_, and four combinations of *k*_1_ and *k*_2_ that sum to *n* = 3: *k*_1_ = 3 and *k*_2_ = 0; *k*_1_ = 2 and *k*_2_ = 1; *k*_1_ = 1 and *k*_2_ = 2; and *k*_1_ = 0 and *k*_2_ = 3. At the moment we assume that demonstrator formation is random. *X_i_* specifies the probability of adoption of trait *i* for that set of *k* values, and incorporates the conformity parameter *a*. If *k_max_* is the maximum *k* in a combination, and *π* is the number of traits that have *k* = *k_max_* (so when *π* = 1 there is a single most-common trait, and when *π* > 1 there are more than one joint-most-common traits), then:

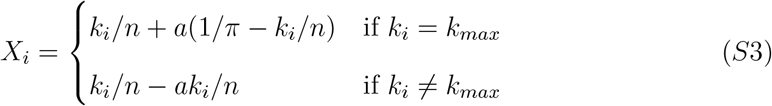

To see how these equations work, Table S2 shows how to calculate the frequency 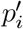 of trait *i* = 1 following conformity given two traits, 1 and 2 (*s* = 2) and three demonstrators (*n* = 3). The first three columns ‘Dem 1-3’ contain all possible combinations of traits 1 and 2 among the three demonstrators. *k*_1_ and *k*_2_ are the number of copies of trait 1 and 2 respectively in that row’s demonstrators. ‘Coef’ contains the number of each combination of demonstrator traits ignoring order, and is the multinomial coefficient. The column *P* (*forming*) gives the probability of that combination of demonstrators forming. This is given by the standard multinomial expression, i.e. the product of each trait frequency (*p* and 1 − *p*, given that there are only two traits) raised to the powers *k*_1_ and *k*_2_ respectively.

**Table S2:**
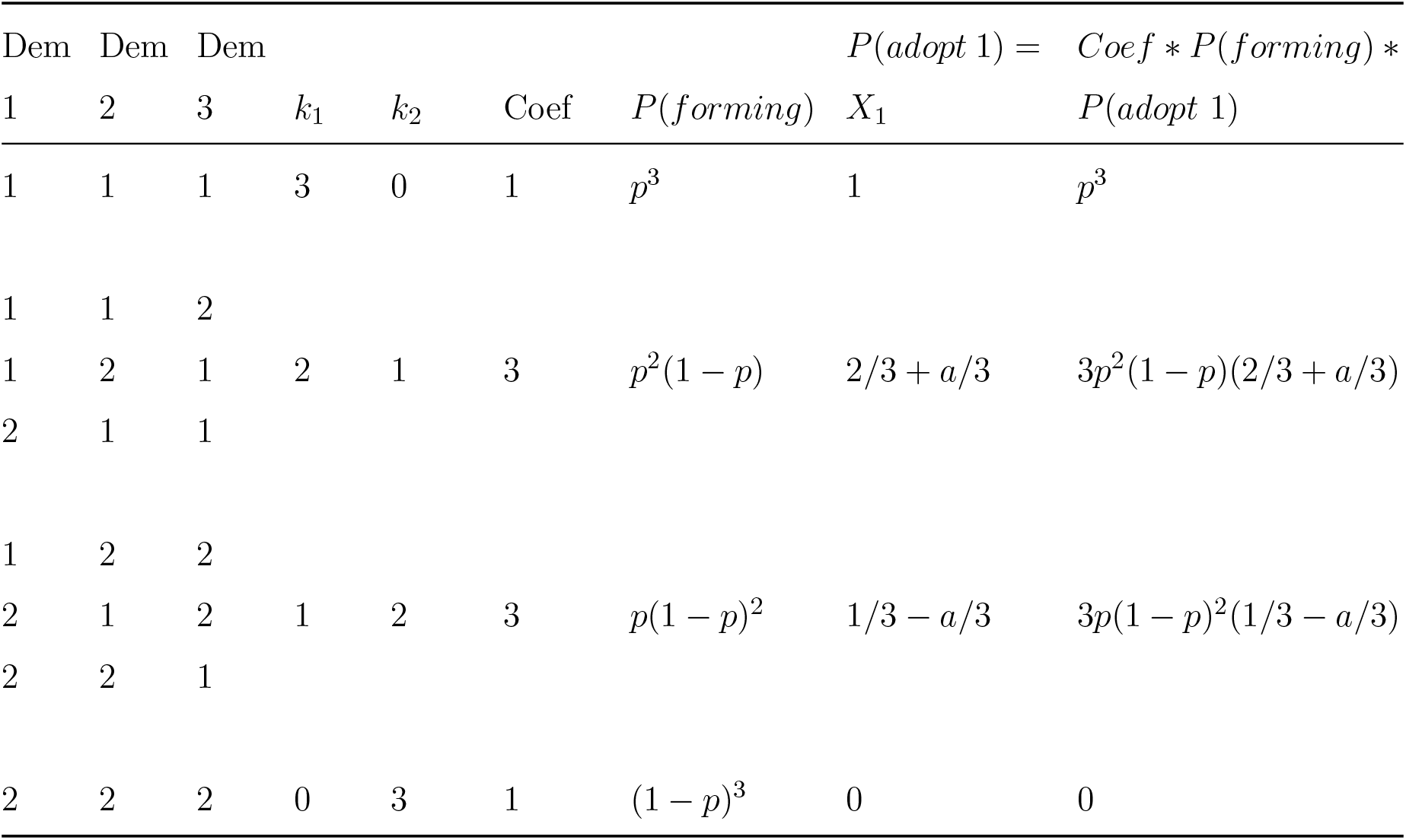
Example frequency table for calculating the frequency 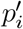 of trait 1 following conformity given two traits, 1 and 2 (*s* = 2) and three demonstrators (*n* = 3) See text for details. Dem=demonstrator, Coef=Coefficient, P(forming)=probability of that demonstrator combination forming randomly, P(adopt 1)=probability of adopting trait 1

*P*(*adopt* 1) gives the probability of that row’s demonstrator trait combination resulting in the observer adopting trait 1, incorporating the conformity parameter *a*. This is *X_i_* in equation S2 and is given in equation S3. In the first row/combination, *k*_1_ = 3 = *k_max_* = *n* and *π* = 1, so *X_i_* = *k*_1_/*n* + *a*(1/*π* − *k_i_*/*n*) = 1 + 0 = 1. For the next three demonstrator combinations, *k*_1_ = 2 = *k_max_* and *π* = 1, so *X_i_* = *k*_1_/*n* + *a*(1/*π* − *k_i_*/*n*) = 2/3 + *a*(1 − 2/3) = 2/3 + *a*/3. And so on for the other demonstrator combinations. The final column contains the product of the coefficient, *P* (*forming*) and *P* (*adopt* 1). The sum of this final column gives the frequency of trait 1 in the next generation, 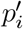:

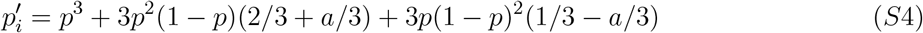

which reduces to equation S1 because *s* = 2 and *n* = 3, as in Boyd and Richerson’s [10] original formulation. When s > 2 or *n* > 3 the resulting recursion will not reduce to equation S1, but is derived in the same way from equation S2. In the case of *s* = 2, then the updated frequency of the other trait will equal 1 − *p*; when *s* > 2 then *s* − 1 traits need to be calculated using Equation S2.

Note that some combinations of *n >* 3 demonstrators will have more than one maximum *k*, e.g. for the trait combination {1,1,2,2,3} then *k*_1_ = 2, *k*_2_ = 2 and *k*_3_ = 1, so *k*_1_ and *k*_2_ are both *k*_max_. In such cases equation S3 also applies, to increase the frequency of both most-common traits equally by the amount that the minority traits are decreased.

Fig S3 shows the effect of increasing the number of demonstrators (*n* > 3) and adding more than two traits (*s* > 2) to the change in trait frequencies over time, with no assortation (*r* = 0). Increasing *n* increases the speed with which majority traits go to fixation. Increasing *s* causes the initially most-common trait to go to fixation, and all other traits to be eliminated.

**Figure S3:**
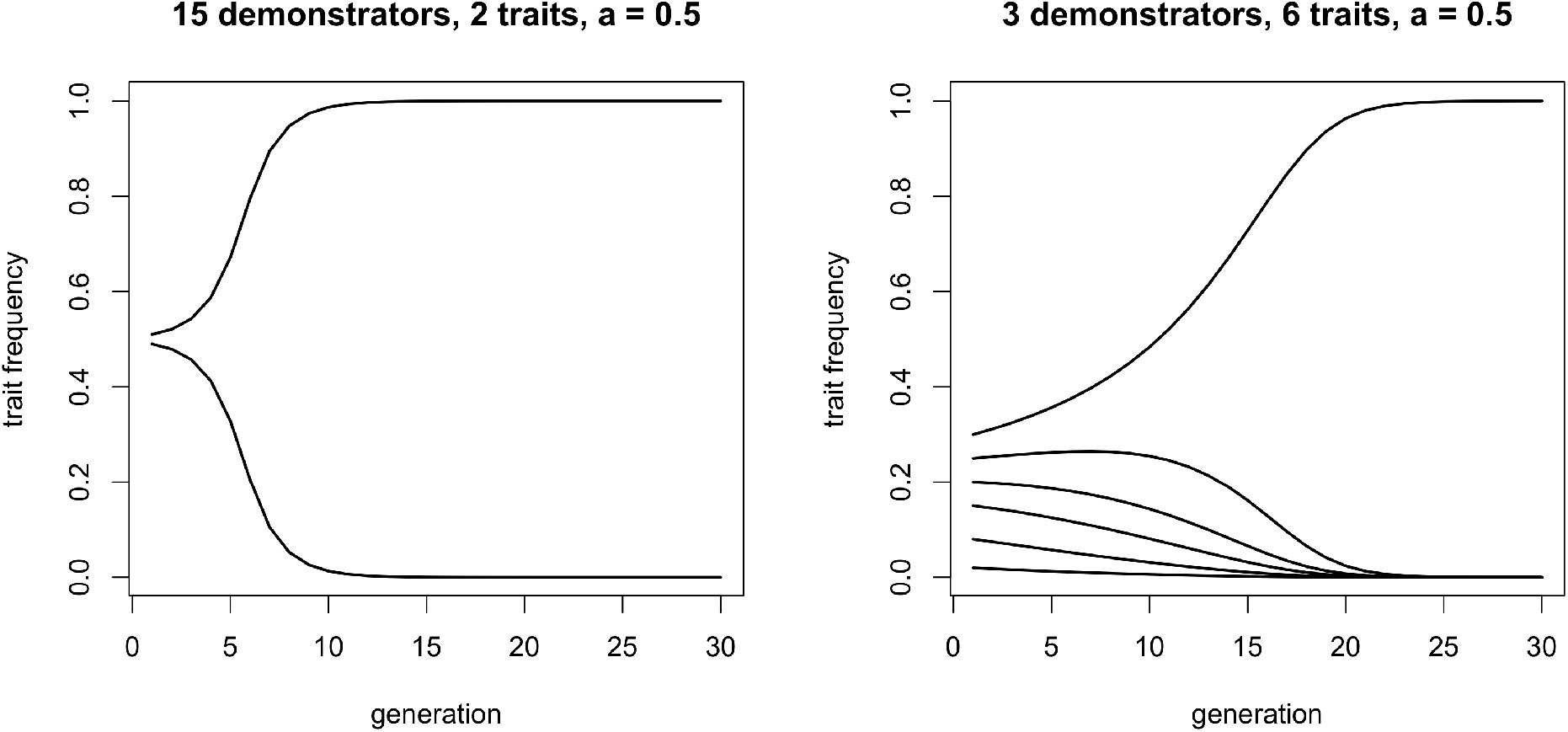
The effect of conformity on trait frequencies, for more than three demonstrators (left) and more than two traits (right). Increasing the number of demonstrators increases the strength of conformity (compare with Figure S2, left panel). For more than two traits, whichever trait is initially most frequent goes to fixation, even if this frequency is initially less than 0.5 (here initial trait frequencies were 0.3, 0.25, 0.2, 0.15, 0.08, and 0.02). Other parameters: *r*=0

Fig S4 shows the probability of adopting a trait given different frequencies of that trait in the population, which have become defining images of conformity in the cultural evolution literature. I show only the case of *s* = 2 for ease of interpretation. The left panel shows that a non-zero conformity parameter *a* generates S-curves, such that when the trait is common (its frequency is greater than 0.5) then the probability of adoption is exaggerated, and when the trait is uncommon (less than 0.5) then the probability of adoption is decreased, relative to the dotted line which shows unbiased, non-conformist transmission. Larger values of *a* generate stronger conformity curves. The right panel shows that increasing *n* also increases the strength of conformity, keeping *a* constant.

**Figure S4:**
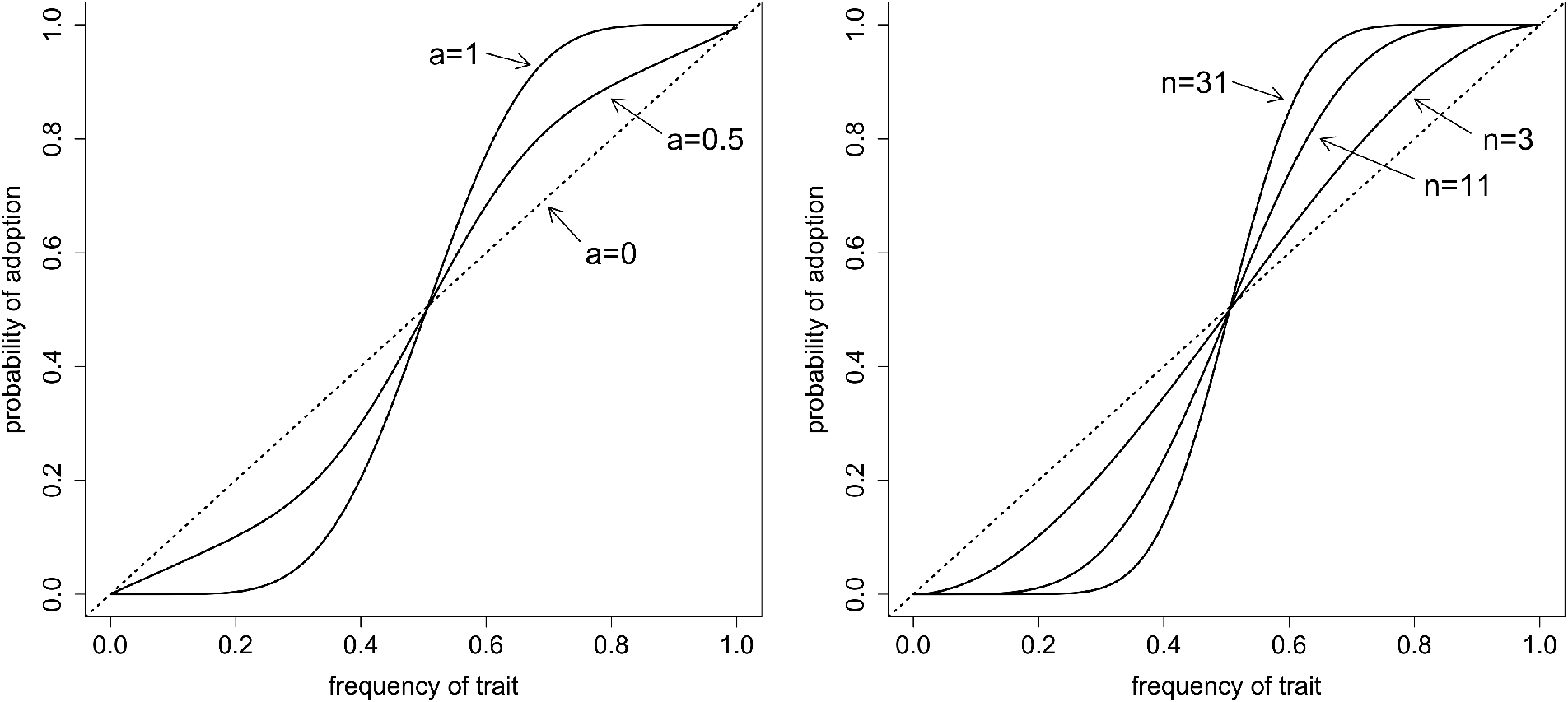
How conformity affects trait adoption. The y-axis shows the probability of adopting a trait as a function of that trait’s frequency in the population. Here we assume only two traits (*s*=2). The dotted line shows unbiased, non-conformist transmission: the probability of adoption is exactly equal to the frequency in the population. The left panel shows different values of *a* for constant *n* (*n*=15). The right panel shows different values of *n* for constant *a* (*a*=1). Other parameters: *r*=0.

We now add non-random formation of demonstrators, or assortation. Boyd and Richerson [10] also implemented assortation, but they restricted its effect to a correlation *r* between two of three demonstrators. Here I wish to add a general assortation effect across all *n* demonstrators, where *n* ≥ 3. The simplest case would be where a fraction 1−*r* of demonstrator combinations form randomly as specified in Equation S2, and a fraction *r* form culturally homogenous sets of demonstrators who all possess the same cultural trait. In Table S2, this would be the first row (all demonstrators have trait 1) and last row (all demonstrators have trait 2). If we are interested in how *p_i_* changes, then only one of these combinations will result in a change in *p_i_* (the one where *k_i_* = *n*) due to the *X_i_* term. Assuming learners must possess the same trait *p_i_* as the homogenous set of demonstrators, then a fraction *p_i_* of individuals will learn from homogenous sets. Homogeneous sets always produce the same trait (again due to the *X_i_* term), and so result in the same frequency of *p_i_* as before transmission. In Equation S5, this is reflected in the *rp_i_* term:

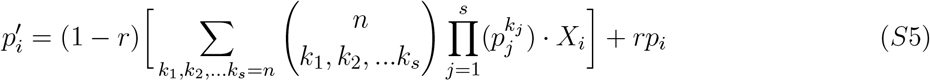

It is clear from Equation S5 that as *r* increases, the effect of conformity via *X_i_* becomes weaker.

### Model 2 fitness plots and assumptions

The fitness functions given in the main text are a specific cooperation version of the general fitness functions for coordination games provided in Boyd & Richerson [13]. Fig S4 plots the fitness of cooperators (*w_c_*, blue line) and defectors (w_d_, red line) at different frequencies of cooperators, *p*, equivalent to Fig 1 in [13]. The lines cross at *p*\*, as given by Equation 4 in the main text. This point *p*\*, where the fitness of cooperators and defectors is equal, is an unstable equilibrium for payoff-biased within-group social learning (as determined by *L*). To ensure that fitnesses are always positive, I assume that *b* > *c*, *b* > *v* and *c* + *u* < 1.

**Figure S5:**
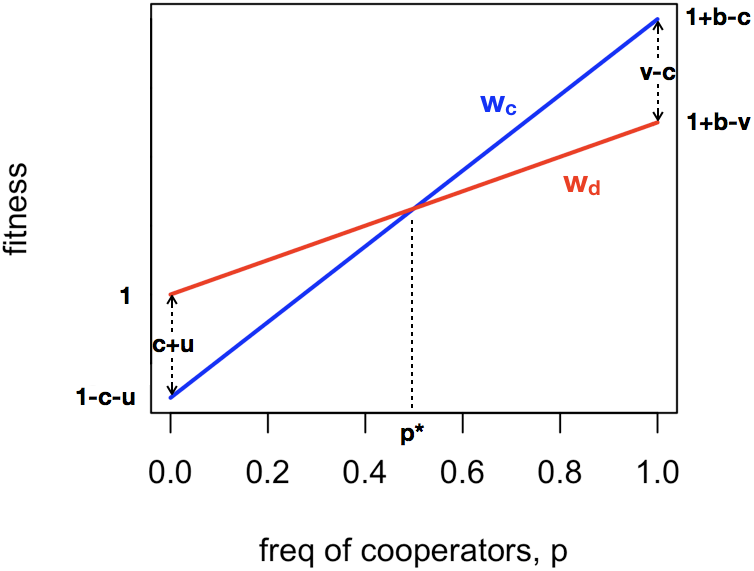
Fitnesses of cooperators (*w_c_*, blue line) and defectors (*w_d_*, red line) at different frequencies of cooperators, *p*. Annotations show how fitness parameters affect these functions, and *p*\* indicates where the lines cross.

The fitness parameter values were chosen in Fig S5 so that *p*\* = 0.5 and the basin of attraction within which cooperators have higher fitness than defectors (*w_c_* > *w_d_* or *p* > *p*\*) and the basin of attraction within which defectors have higher fitness than cooperators (*w_d_* > *w_c_* or *p* < *p*\*) are equal. Other fitness parameter values give different values of *p*\* and alter the relative sizes of these basins of attraction. The easiest way of doing this is varying *v*, the punishment cost borne by defectors. Fig S6 shows fitness plots for three different values of v, in the centre with equal basins of attraction (as in Fig S5), on the right where cooperators have a larger basin of attraction, and on the left where defectors have a larger basin of attraction. This has consequences for the action of payoff-biased within-group social learning (L) by changing the unstable equilibrium point.

**Figure S6:**
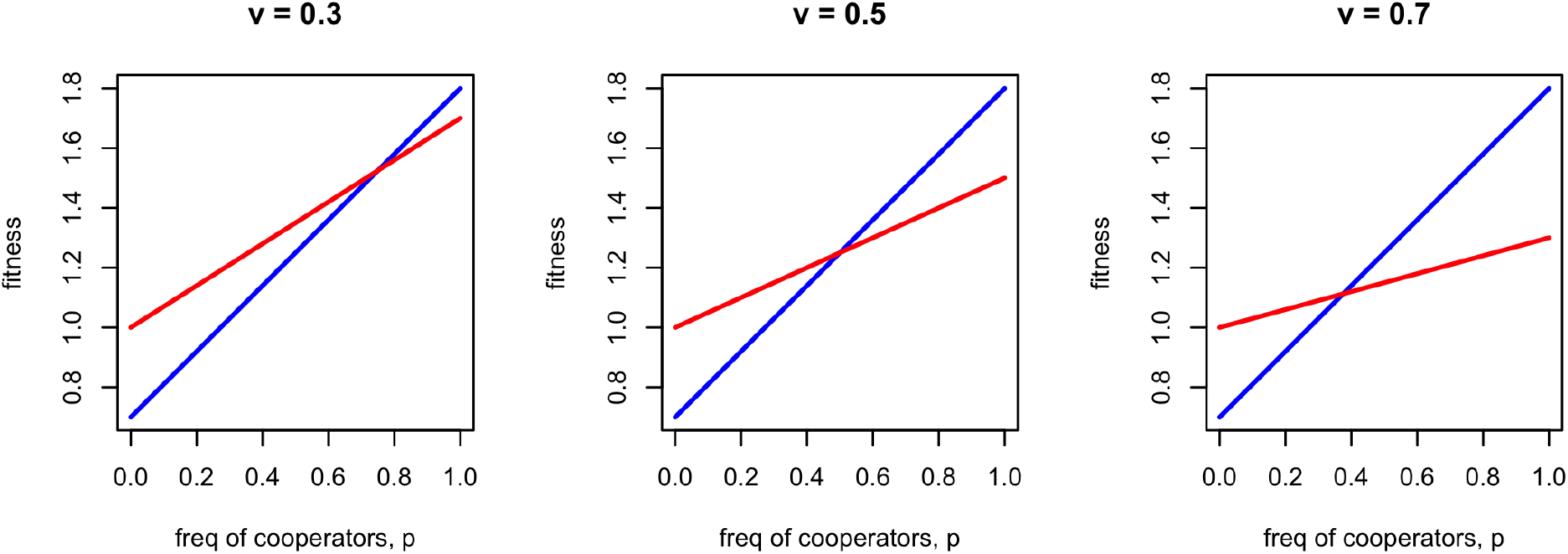
Fitnesses plots for three different values of *v*, which determines the cost of being punished for defectors. When *v* is large, cooperators have a larger basin of attraction. When *v* is small, defectors have a larger basin of attraction. Other parameters: *b*=1, *c*=0.2, *u*=0.1.

Equation S6 gives the mean population fitness *W*. Fig S7 plots this quadratic function of *p*.

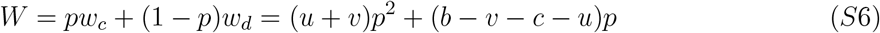

To keep the model simple I assume that *W* is always greater than 1, in other words, a population entirely composed of defectors (*p* = 0) always has lower fitness than any population containing any cooperators (*p* > 0). In terms of Fig S7, this would mean that the green *W* line never drops below *W* = 1. To ensure this, I assume that (*b*−*c*) > (*u*+*v*). Finally, the parameter *μ* in Equation 5 ensures that the weighted migration rate *m*(*W* − 1) never exceeds the baseline migration rate *m*. This is done by setting *μ* to be the reciprocal of the maximum value of *W*−1 which occurs at *p* = 1, so *μ* = 1/(*b*−*c*). Similarly, *γ* in Equation 6 ensures that the change in *p* due to payoff-biased social learning always scales from 0 to *L*. From Fig S5, the maximum absolute fitness difference between cooperators and defectors (*w_c_* − *w_d_*) is either *v* − *c* or *c* + *u*, whichever is larger, so *γ* = 1/[*max*(*v* − *c*, *c* + *u*)].

**Figure S7:**
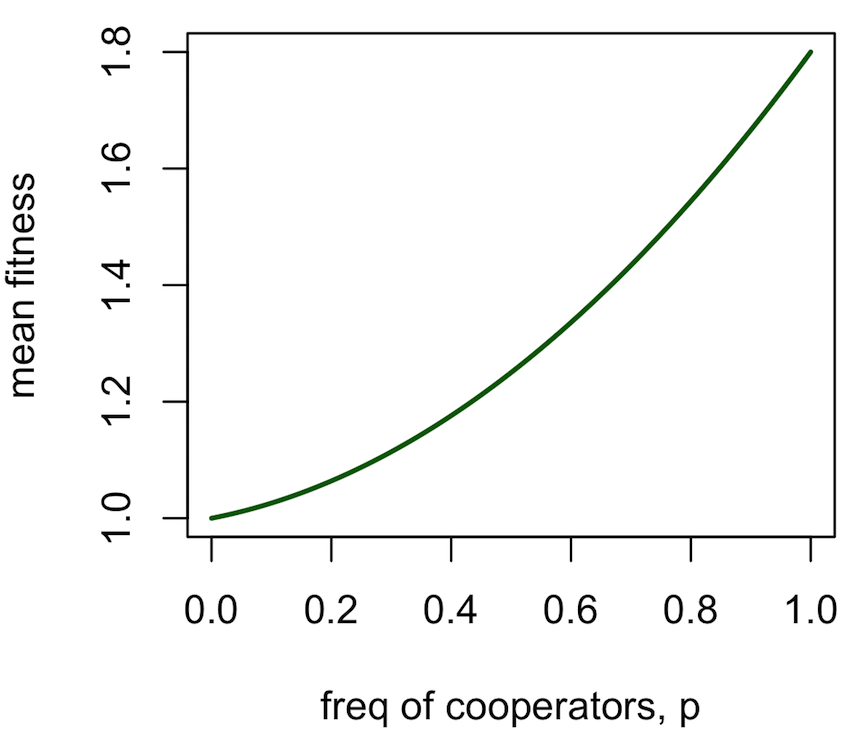
Mean sub-population fitness *W* as a function of *p*, as given by Equation S5. Parameters: *b*=1, *c*=0.2, *u*=0.1, *v*=0.5.

## S1 Results

### Individual-based version of Model 1

In order to verify the recursion-based models that track trait frequencies, I created analogous individual-based models in which individuals and their traits are explicitly simulated. These individual-based models have the same assumption of *s* sub-populations and *s* traits. Now, however, we specify the number of individuals in each sub-population, *N*. This is different to *n*, which is the number of demonstrators sampled during conformist acculturation. *n* is also implemented in the individual-based models, so individuals pick *n* demonstrators from the *N* members of their sub-population (so *n* ≤ *N*). The individual-based simulations start with the same complete between-group structure as the recursion models, i.e. all *N* individuals in sub-population 1 have trait 1, all *N* individuals in sub-population 2 have trait 2, etc. *F_ST_* is calculated in the same way as for the recursion-based model after calculating overall trait frequencies from individuals’ traits. Migration occurs in the same way as the island model. Every time-step, each individual moves to a common migrant pool with probability *m*, and then randomly disperses across the newly vacant spots ignoring sub-population structure. Conformist acculturation is implemented slightly differently. Following all migration, each individual chooses *n* demonstrators from within their sub-population, and with probability *a* adopts the most common trait among those *n* demonstrators. Otherwise they retain their existing trait. This is conceptually the same as the recursion-based conformity described above, but without the multinomial theorem implementation, thus providing a conceptual replication of conformity as implemented in the main text. For each of the *n* chosen demonstrators, with probability 1 − *r* that demonstrator is chosen randomly from the focal individual’s sub-population. With probability *r* that demonstrator has the same trait as the focal individual. Consequently, when assortation parameter *r* =1, then individuals only ever learn from demonstrators with the same trait as themselves. Parameter definitions for the individual-based model are given in Table S3. Full code of all models is available in https://github.com/amesoudi/migrationmodels.

**Table S3:** Parameter definitions for Model 1 individual-based simulations

| Parameter | Definition |
| --- | --- |
| $s$ | The number of sub-populations, and also the number of alternative cultural trait values |
| $m$ | Migration rate: the probability in one time-step that an individual moves to a randomly chosen sub-population (equivalently, the proportion of the population that moves in one time-step) |
| $n$ | The number of demonstrators from whom individuals learn during acculturation |
| $a$ | Acculturation rate: the probability that an individual adopts the most-common trait among $n$ demonstrators chosen from their sub-population, as opposed to retaining their existing trait. When $a = 1$, there is 100% chance of copying the most-common trait (if there is one). |
| $N$ | The number of individuals in each sub-population (giving $Ns$ individuals in the entire population) |
| $r$ | The probability that a demonstrator is chosen who has the same trait as the focal individual, rather than chosen at random. |

Figs S8, S9 and S10 show the individual-based model results equivalent to the recursion-based model results shown in Figs 1, 2 and 3 respectively. There is very little difference between the two models, despite their different implementation, increasing our confidence in the robustness of the conclusions. There is a slight tendency for *F_ST_* to be lower in the individual-based model than the recursion-based model, especially when diversity is low (e.g. *a* = 0.6 in Fig S8C). This may be due to drift which operates only in the finite populations of the individual-based models. If a trait is lost due to drift, then diversity will be reduced. Fig S11 repeats Figs 2 and S10 but with larger *n* than is computationally feasible with the recursion-based model. Here we can see that increasing *n* above 13 does not change the dynamics, except at very low values of *a* when migration is very high (Fig S11C).

**Figure S8:**
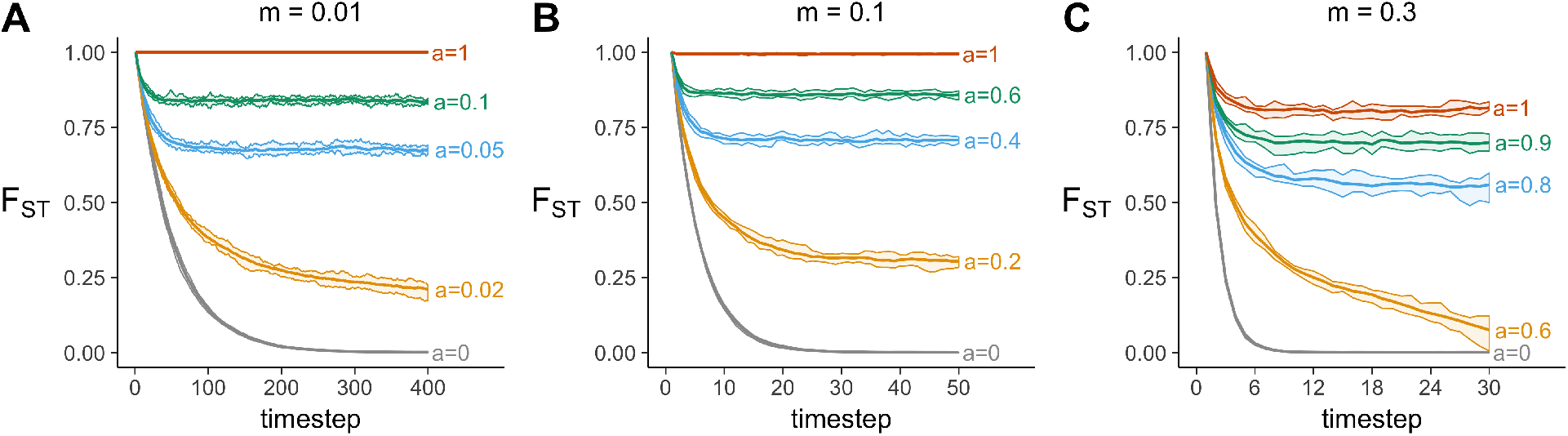
Time series showing changes in *F_ST_* over time for (A) a low migration rate *m*=0.01, (B) a moderate migration rate *m*=0.1, and (C) a high migration rate *m*=0.3, at varying strengths of acculturation, *a*, in the individual-based model. Other parameters: *s*=5, *n*=5, *N*=1000; results are the average of 10 independent simulation runs.

**Figure S9:**
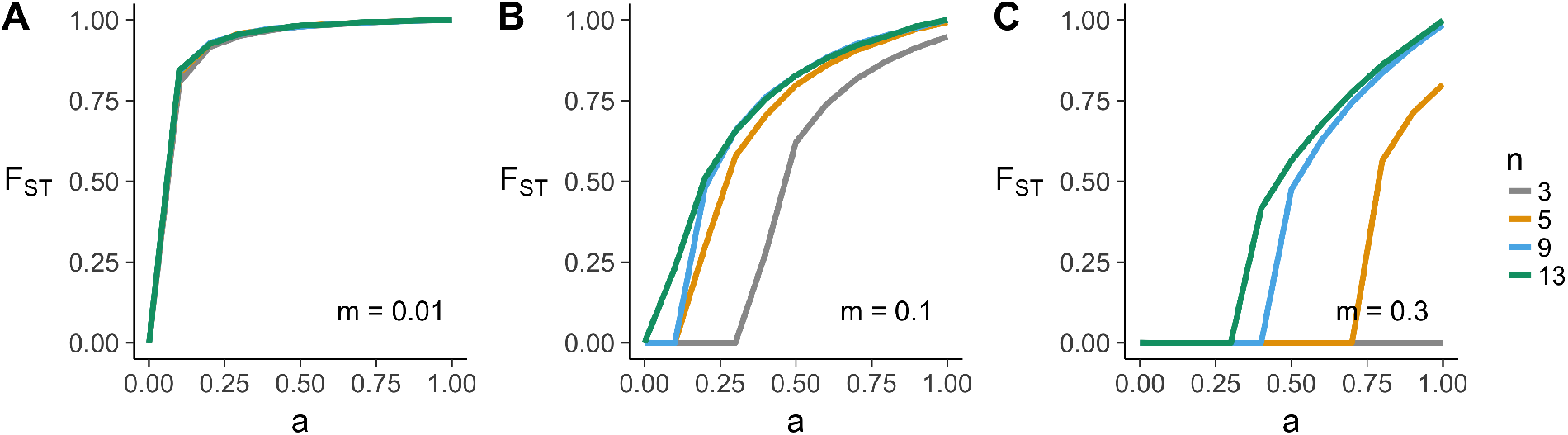
The relationship between *a* and *F_ST_* at three different migration rates, and different values of n, for the individual-based model. Other parameters: *s*=5, 500 timesteps, *N*=1000; results are the average of 10 independent simulation runs.

**Figure S10:**
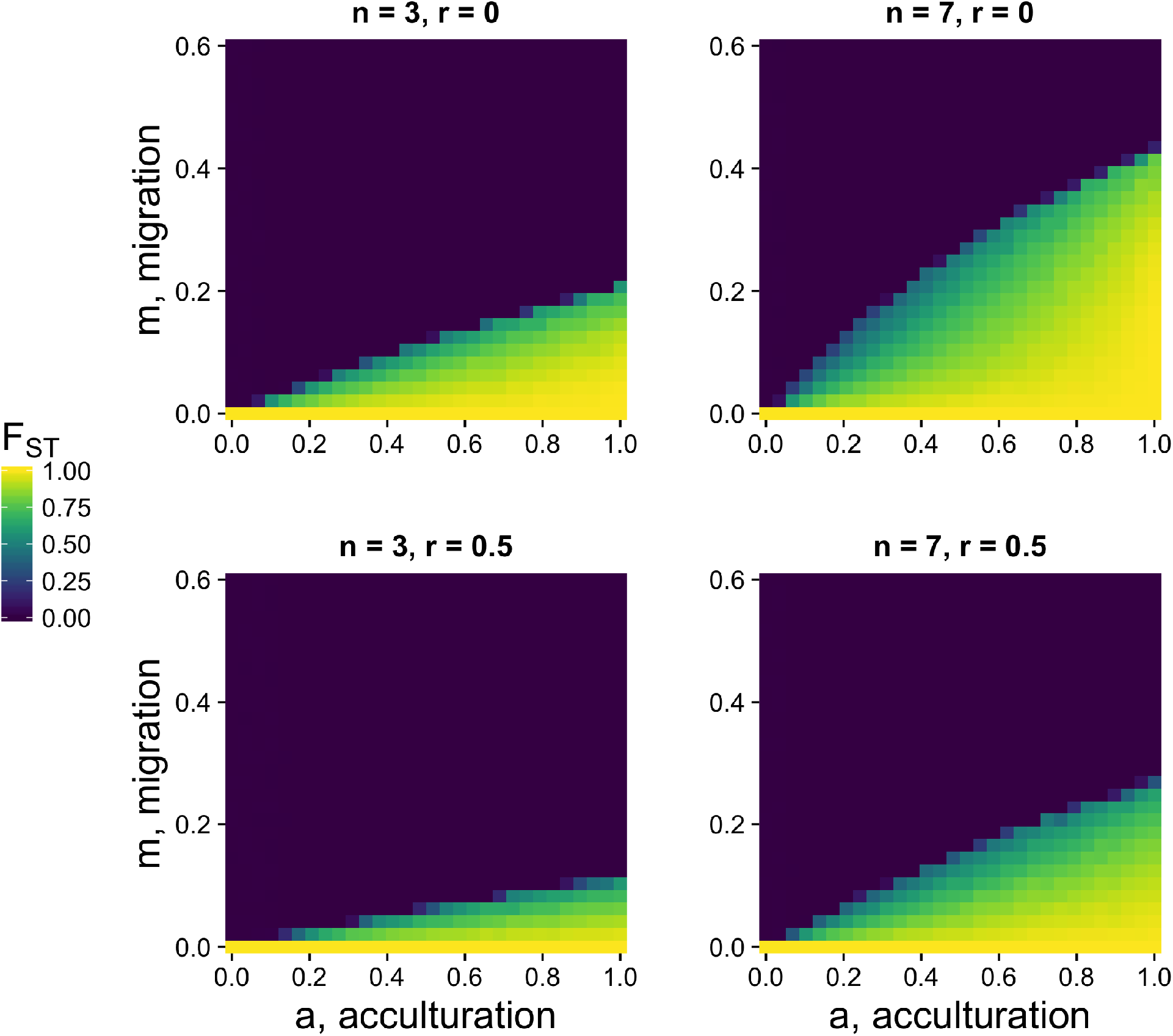
Heatmap showing *F_ST_* for varying acculturation rates, *a*, and migration rates, *m*, separately for three different values of n, the number of demonstrators, for the individual-based model. Other parameters: *s*=5, *N*=1000, 500 timesteps; results are the average of 10 independent simulation runs.

**Figure S11:**
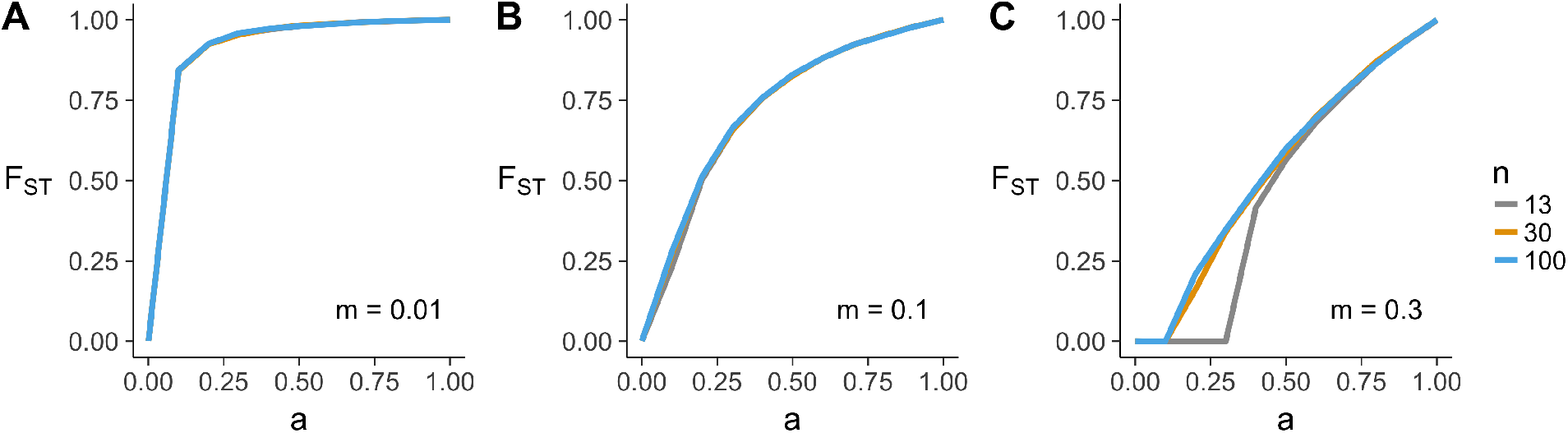
The relationship between *a* and *F_ST_* at three different migration rates, for three large values of *n*, for the individual-based model. Other parameters: *s*=5, 500 timesteps, *N*=1000; results are the average of 10 independent simulation runs.

